# The Oncoprotein BCL6 Enables Cancer Cells to Evade Genotoxic Stress

**DOI:** 10.1101/2021.06.15.448559

**Authors:** Yanan Liu, Juanjuan Feng, Kun Yuan, Yue Lu, Kun Li, Jiawei Guo, Chengbin Ma, Jing Chen, Xiufeng Pang

## Abstract

Genotoxic agents remain the mainstay of cancer treatment. Unfortunately, the clinical benefits are often countered by a rapid tumor adaptive response. Here, we report that the oncoprotein B cell lymphoma 6 (BCL6) is a core component that confers tumor adaptive resistance to genotoxic stress. Multiple genotoxic agents promoted BCL6 transactivation, which was positively correlated with a weakened therapeutic efficacy and a worse clinical outcome. Mechanistically, we discovered that treatment with the genotoxic agent etoposide led to the transcriptional reprogramming of multiple pro-inflammatory cytokines, among which the interferon-α and interferon-γ responses were selectively and substantially enriched in resistant cells. Our results further revealed that the activation of interferon/signal transducer and activator of transcription 1 axis directly upregulated BCL6 expression. The increased expression of BCL6 further repressed the tumor suppressor PTEN and consequently enabled resistant cancer cell survival. Accordingly, targeted inhibition of BCL6 remarkably enhanced etoposide-triggered DNA damage and apoptosis both *in vitro* and *in vivo.* Our findings highlight the importance of BCL6 signaling in conquering tumor tolerance to genotoxic stress, further establishing a rationale for a combined approach with genotoxic agents and BCL6-targeted therapy.

## Introduction

Genome instability is the major hallmark of chronic proliferating tumors (Hanahan & Weinberg, 2011; Murai, Thomas, Miettinen, & Pommier, 2019). Conventional genotoxic chemotherapy (e.g., topoisomerase II inhibitors, cisplatin, carboplatin) that introduce DNA damage lesions, devastate genomic integrity and activate pro-apoptotic pathways, are employed as the standard first-line treatment for a wide array of solid malignancies (Cheung-Ong, Giaever, & Nislow, 2013). Despite initial therapeutic success, intrinsic resistance or rapid adaptive resistance in cancer cells is a major hurdle, hampering the clinical efficacy of these agents (O’Grady et al., 2014; Stebbing et al., 2018; Trinh, Ko, Barengo, Lin, & Naora, 2013). Chemoresistance occurs due to complex reasons, such as an increased DNA damage repair capacity, activation of pro-survival pathways, and defects in caspase activity (Poth et al., 2010; Stebbing et al., 2018). While several signaling effectors have been identified as predictive markers, such as *ABCA1* (Koh et al., 2019) and *MAST1* (Jin et al., 2018), in tumor tolerance to genotoxic agents, the majority of these studies lacked either an evaluation of the clinical correlation or an explanation for how these effectors mediate pro-survival signals in the presence of genotoxic stress.

The transcriptional repressor B cell lymphoma 6 (BCL6) has emerged as a critical therapeutic target in diffuse large B-cell lymphomas (Parekh, Prive, & Melnick, 2008). Increasing evidences indicate that BCL6 plays an oncogenic role in several human hematopoietic malignancies and solid tumors (Beguelin et al., 2016; Cardenas et al., 2017; Deb et al., 2017). BCL6 binds and represses different target genes to drive tumorigenesis in a cell context-dependent manner (Ci et al., 2009). The constitutive expression of BCL6 sustains the lymphoma phenotype and promotes glioblastoma through transcriptional repression of the DNA damage sensor ATR (Ranuncolo et al., 2007) and the p53 pathway (Xu et al., 2017), respectively. According to data derived from The Cancer Genome Atlas (TCGA), the BCL6 locus is also predominantly amplified in primary breast cancer and is correlated with a worse prognosis (Walker et al., 2015). Recently, small molecular inhibitors that target the interaction between BCL6 and its co-repressors or that trigger BCL6 degradation effectively restored BCL6 target gene expression and impeded tumor growth (Cardenas et al., 2016; Cheng et al., 2018; Slabicki et al., 2020).

The properties of BCL6 as a therapeutic target originate from its normal function in sustaining the proliferative and the phenotype of stress-tolerant germinal center B cells (Phan, Saito, Kitagawa, Means, & Dalla-Favera, 2007). BCL6 allows B cells to evade ATR-mediated checkpoints and tolerate exogenous DNA damage by repressing the cell cycle checkpoint genes *CDKN1A*, *CDKN1B*, and *CDKN2B*, and the DNA damage sensing genes *TP53*, *CHEK1*, and *ATR* (Basso et al., 2010; Cardenas et al., 2017; Phan, Saito, Basso, Niu, & Dalla-Favera, 2005). When genotoxic stress is accumulated to some extent, BCL6 is phosphorylated by the DNA damage sensor ATM kinase and degraded through the ubiquitin proteasome system, whereby the germinal center reaction is terminated (Phan et al., 2007). The critical functions exerted by BCL6 during normal B cell development could be hijacked by malignant transformation, thereby leading to lymphoma (Basso & Dalla-Favera, 2012). Recent studies have suggested that BCL6 is involved in stress tolerance and drug responses. In detail, BCL6 can be activated by heat shock factor 1 to tolerate heat stress (Fernando et al., 2019). The aberrant expression of BCL6 can be provoked in leukemia cells in response to the tyrosine kinase inhibitor imatinib (Duy et al., 2011). Our recent work additionally revealed that an increased expression of BCL6 largely contributes to the resistance of *KRAS*-mutant lung cancer clinical BET inhibitors (Guo et al., 2021). Given the fact that BCL6 plays an emerging role in DNA damage tolerance and drug responses, we hypothesized that BCL6 might drive cancer cell resistance to genotoxic agents.

Here, we report that the proto-oncogene BCL6 is a central component of the resistance pathway in tumor response to genotoxic agents. We observed a striking association between the activation of pro-inflammatory signals and BCL6 induction in chemoresistant cancer cells. The tumor suppressor PTEN is further characterized as a functional target gene of BCL6. Importantly, addition of BCL6-targeted therapy to the genotoxic agent etoposide markedly restored the sensitivity of cancer cells to etoposide *in vitro* and *in vivo*. Overall, our findings establish a rationale for targeting BCL6 to conquer resistance to genotoxic stress in solid tumors.

## Results

### Genotoxic agents promote BCL6 transcription

While genotoxic agents have become the mainstay of clinical cancer treatments (Fillmore et al., 2015; Nitiss, 2009), many patients show a poor response to these drugs due to the emergence of a tumor rapid adaptive response (Wijdeven et al., 2015). To gain a comprehensive understanding of chemoresistance mechanisms, we initially measured the half inhibitory concentrations (IC_50_s) of etoposide and doxorubicin, two well-validated topoisomerase II inhibitors for clinical use, in a panel of 22 cancer cell lines derived from four types of solid tumors, including lung, pancreatic, colorectal, and ovarian carcinomas. Some cell lines displayed apparent resistance to etoposide at doses up to 30 μM (**Figure 1A**) or to doxorubicin at doses up to 0.6 μM (**Figure 1-figure supplement 1A**), while the remaining cell lines showed a gradient of sensitivity to them. The concentrations of 30 μM and 0.6 μM were chosen to define the resistance of multiple cancer cell lines to etoposide and doxorubicin, respectively, as these are the highest achievable concentration in the plasma of patients, which are likely to be clinically relevant (Kaul et al., 1995; Palle et al., 2006).

**Figure 1.**
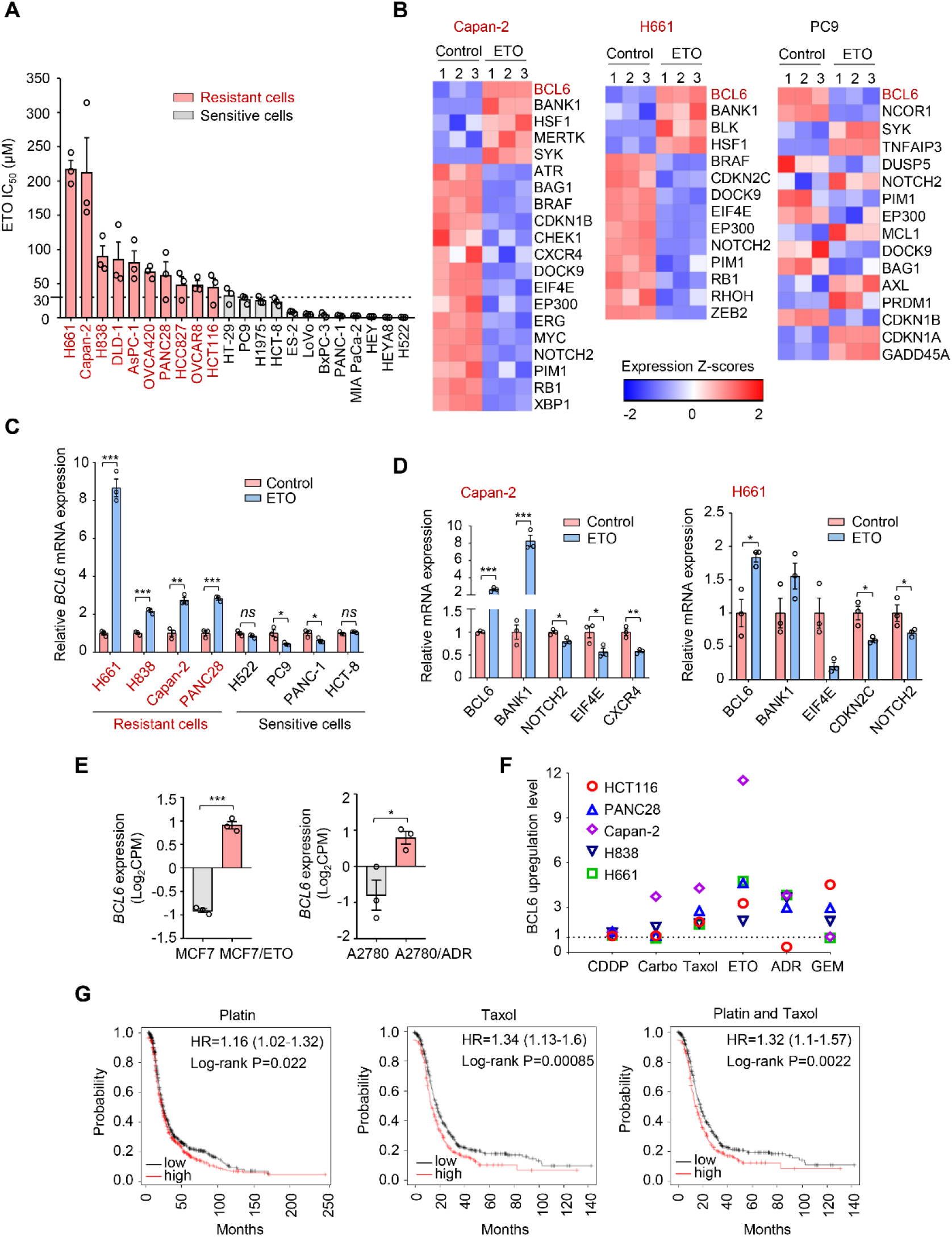
Genotoxic agents promote BCL6 expression. (**A**) Cell sensitivity to etoposide (ETO). Cancer cells were treated with etoposide at gradient concentrations for 48 h. IC_50_s were measured using Sulforhodamine B (SRB) assays. Values are expressed as mean ± SEM of three independent experiments. ETO-resistant cell lines are marked in red. Cell sensitivity to doxorubicin (ADR) was also examined (see **Figure 1-figure supplement 1A**). (**B**) Heat map illustrating expression of BCL6 target genes in Capan-2, H661 and PC9 cell lines. Cells were treated with etoposide at their respective 1/2 IC_50_s for 24 h. mRNA was isolated from treated cells and sequenced. Z-scores were calculated based on counts of exon model per million mapped reads. BCL6 target genes were identified by a cutoff of *P* < 0.05, *n* = 3. (**C**) BCL6 mRNA expression in ETO-resistant and -sensitive cells. Cells were treated with etoposide at their respective 1/2 IC_50_s for 24 h. QPCR assays were subsequently performed. Values are expressed as mean ± SEM of three independent experiments. \**P* < 0.05, \*\**P* < 0.01 and \*\*\**P* < 0.001, unpaired, two tailed *t*-test. ETO-resistant cell lines are marked in red. (**D**) Validation of differentially expressed target genes of BCL6 in Capan-2 and H661 cells using qPCR analysis. Values are expressed as mean ± SEM of three independent experiments. \**P* < 0.05, \*\**P* < 0.01 and \*\*\**P* < 0.001, unpaired, two tailed *t*-test. (**E**) Normalized BCL6 mRNA expression levels in MCF7 and MCF7/ETO (required ETO-resistant MCF7), or A2780 and A2780/ADR (required ADR-resistant A2780). Values are expressed as mean ± SEM of three independent experiments. \**P* < 0.05, \*\*\**P* < 0.001, unpaired, two tailed *t*-test. (**F**) BCL6 protein expression levels in different cancer cell lines in response to various genotoxic agents. Cells were treated with indicated genotoxic agents for 24 h. BCL6 protein expression levels were detected and normalized to GAPDH expression using immunoblotting analysis. Representative images related to **Figure 1-figure supplement 1B**. The ratio of genotoxic agent-treated groups to the control group was calculated. CDDP, cisplatin; Carbo, carboplatin; GEM, gemcitabine. (**G**) Kaplan-Meier curves of ovarian cancer patients treated with cisplatin, taxol or both drugs. The curves were stratified by BCL6 (215990_s_at) expression. The following source data and figure supplements are available for figure 1: **Figure 1-Source data 1.** Genotoxic agents promote BCL6 expression. **Figure 1-figure supplement 1.** Genotoxic agents promote BCL6 expression.

To decipher the mechanisms of tumor resistance to genotoxic therapy, we first performed RNA sequencing in etoposide-resistant cells (Capan-2 and H661) and etoposide-sensitive cells (PC9) in the presence or absence of etoposide. An in-depth comparison of the transcriptome was conducted to describe the transcriptional programs that were responsive to etoposide in sensitive cells but remained recalcitrant to treatment in a resistant population. By analyzing the significantly differentially expressed genes, we strikingly found that etoposide treatment triggered a remarkable increase in BCL6 expression in etoposide-resistant Capan-2 and H661 cells, but not in etoposide-sensitive PC9 cells (**Figure 1B**). Given that BCL6 signaling gene sets have not been fully defined in solid tumors, several studies have focused on BCL6 transcriptional program (Ci et al., 2009; Green et al., 2014). In addition to the well-known BCL6 target genes or co-repressors in germinal centers and multiple tumors, such as *BMI1*, *EIF4E*, *NOTCH2* and *BCL2* (Basso et al., 2010; Cerchietti et al., 2010; Ci et al., 2009; Dupont et al., 2016; Valls et al., 2017), several other genes directly regulated by BCL6 have been recently identified using chromatin immunoprecipitation followed by sequencing, including BCL6-activated genes (e.g., *SYK*, *BANK1*, *BLK*, and *MERTK*) or BCL6-repressed genes (e.g., *CDKN2C*, *CDKN1B*, *RB1*, and *PTPRO*) (Geng et al., 2015). We used comparative BCL6 target gene selection to identify the genes that were differentially expressed between resistant and sensitive cells in the presence or absence of etoposide. Our data revealed that the BCL6 transcriptional program was dramatically affected by etoposide in treated Capan-2 and H661 cells, but not in treated PC9 cells (**Figure 1B**). We further verified the specificity of BCL6 increase in other etoposide-resistant cell lines (**Figure 1C**) and the effects of etoposide on BCL6 target gene expression using qPCR analysis (**Figure 1D**). Given that BCL6 transcription was induced in primary chemoresistant cells, we next tested whether it could be provoked in acquired chemoresistant cells. Therefore, we analyzed published microarray data (Januchowski, Zawierucha, Rucinski, Nowicki, & Zabel, 2014; Moitra et al., 2012), and found that BCL6 upregulation was also observed in acquired resistance process (**Figure 1E**)

To clarify whether the fact that transcriptional induction of BCL6 confers tolerance to genotoxic stress was a general phenomenon, we treated five cell lines with a panel of frontline genotoxic agents (Ettinger et al., 2017; Sandler et al., 2006; Tempero et al., 2017). The results showed that BCL6 was upregulated in response to the majority of these clinical agents (**Figure 1F** and **Figure 1-figure supplement 1B**). In addition, a high expression of BCL6 was associated with a poor progression-free survival in patients who received cisplatin, taxol, or both drugs (**Figure 1G**). These results collectively suggest that an aberrant BCL6 expression might contribute to chemoresistance and is linked to a poor prognosis.

### BCL6 transactivation is correlated with therapy resistance

To further identify whether an increased BCL6 expression was associated with the therapeutic efficacy of genotoxic agents, we first examined BCL6 protein expression in a panel of solid tumor cell lines treated with etoposide or doxorubicin. In agreement with the BCL6 transcription pattern observed in **Figure 1C**, BCL6 protein abundance was dramatically and preferentially induced by etoposide in resistant cells, whereas it was decreased or unchanged in sensitive cells (**Figure 2-figure supplement 1A**). Notably, increased BCL6 protein levels were closely associated with increased etoposide IC_50_ values (**Figure 2A**). Specifically, cells with higher BCL6 protein levels were prone to be more tolerant to etoposide (R^2^ = 0.61, *P* < 0.0001; **Figure 2B**). Similar results were also obtained for doxorubicin (**Figure 2-figure supplement 1B**).

**Figure 2.**
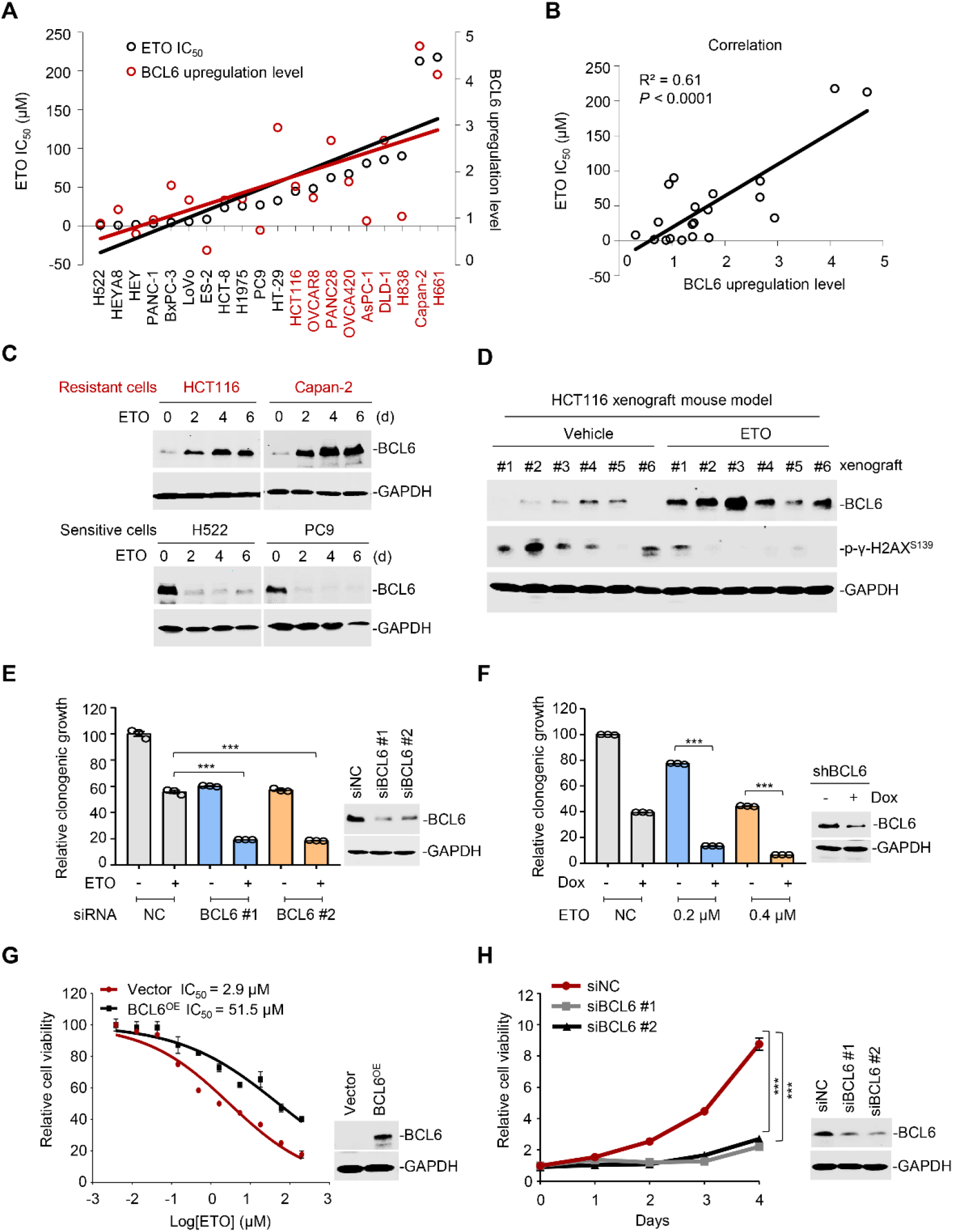
BCL6 transactivation is correlated with therapy resistance. (**A**) Association between BCL6 upregulation with ETO sensitivity in various cancer cell lines. Representative images related to **Figure 2-figure supplement 1A**. Left vertical axis, IC_50_s of etoposide in different cancer cell lines; right vertical axis, relative BCL6 protein levels compared with that of the control group; horizontal axis, cancer cell lines. (**B**) Correlation analysis. Correlation between BCL6 upregulation levels and ETO IC_50_s or ADR IC_50_s (see **Figure 2-figure supplement 1B**). (**C**) Etoposide induced BCL6 protein expression in a time-dependent manner. ETO-resistant or -sensitive cells were treated with etoposide at their respective 1/4 IC_50_s for 2, 4 or 6 days. Cell lysates were collected and probed with specific antibodies using Western blotting assays. ETO-resistant cell lines are marked in red. (**D**) Etoposide increased BCL6 expression and decreased the phosphorylated levels of γ-H2AX (S139) in HCT116 xenografts treated with 10 mg/kg etoposide for 14 days. At the end of the experiment, tumor tissues were isolated and subjected to immunoblotting analysis. Six biologically independent samples of each group are shown. Tumor volume curves and tumor weight are shown in **Figure 2-figure supplement 1C**. (**E**) Relative clonogenic growth of ETO-resistant cells. HCT116 cells were transfected with BCL6 siRNAs or the control siRNA, followed by the treatment of 0.2 μM etoposide for 7 days. The expression of BCL6 was detected by immunoblotting analysis (*right*). Values are expressed as mean ± SEM of three independent experiments by setting the control group as 100%. \*\*\**P* < 0.001, unpaired, two tailed *t*-test. (**F)** Relative clonogenic growth of ETO-resistant cells. HCT116 cells stably transfected with shRNA targeting BCL6 were exposed to etoposide (0.2 or 0.4 μM) with or without doxycycline (Dox) for 7 days. The clonogenic growth were examined. The BCL6 expression levels were detected by an immunoblotting assay (*right*). Values are expressed as mean ± SEM. \*\*\**P* < 0.001, unpaired, two tailed *t*-test. (**G**) BCL6 overexpression decreased the sensitivity of H522 cells to etoposide (*left*). ETO-sensitive H522 cells were transfected with pcDNA3.1-BCL6 or pcDNA3.1 control plasmid, and then treated with etoposide at gradient concentrations for 48 h. The etoposide IC_50_s were detected by SRB assays. BCL6 overexpression efficiency was examined by an immunoblotting assay (*right*). (**H**) Cell viability curves of required doxorubicin-resistant cells in response to BCL6 knockdown. MCF7/ADR cells were transfected with siRNAs targeting BCL6 or the control siRNA. Data are presented as mean ± SEM of six independent experiments by setting the control group as 1. \*\*\**P* < 0.001, unpaired, two tailed *t*-test (*right*). The following source data and figure supplements are available for figure 2: **Figure 2-Source data 1.** BCL6 transactivation is correlated with therapy resistance. **Figure 2-figure supplement 1.** BCL6 upregulation is associated with therapy resistance.

A more detailed observation demonstrated that BCL6 protein expression could also be time-dependently provoked by a long-term exposure of resistant cells to etoposide (**Figure 2C**). This prompted us to examine the responsive role of BCL6 *in vivo*. Therefore, we set up a xenograft mouse model using human HCT116 cells and examined the BCL6 expression shift in xenografts after etoposide treatment. Although etoposide impeded tumor growth at a dose of 10 mg/kg/day (**Figure 2-figure supplement 1C**), the protein level of phosphorylated H2AX, a DNA damage marker (Bonner et al., 2008), was overall decreased in etoposide-treated xenografts compared with that in the vehicle group (**Figure 2D**), implying the emergence of drug resistance. In contrast with the decreased level of phosphorylated H2AX, the BCL6 protein levels in the xenografts were dramatically increased by etoposide, which was consistent with our *in vitro* observations (**Figure 2C**), suggesting that a reciprocal alteration of BCL6 expression is associated with tumor responses to genotoxic agents.

Next, to examine whether BCL6 transactivation affects drug efficacy in resistant cells, we targeted BCL6 using two different small interfering RNAs and found that BCL6 genetic knockdown dramatically attenuated the clonogenic growth of HCT116 cells in the presence of etoposide (**Figure 2E**). In line with these results, inducible knockdown of BCL6 potentiated the killing effects of etoposide (**Figure 2F**). In addition, we overexpressed BCL6 using a lentiviral vector in etoposide-sensitive H522 cells and tested the cytotoxicity of etoposide. As expected, our results showed that BCL6 overexpression increased the etoposide IC_50_ by up to 17-fold (**Figure 2G**). To further investigate the role of BCL6 in adaptive resistance, we introduced siBCL6 into acquired doxorubicin-resistant cells, and found that BCL6 depletion was sufficient to suppress cell proliferation of MCF7/ADR cells (**Figure 2H**). Collectively, these data support the notion that BCL6 confers drug resistance and induces a targetable vulnerability in tumor cells.

### Genotoxic stress activates interferon responses

To further elucidate the mechanisms of BCL6 feedback activation, we conducted gene ontology enrichment analysis on the transcripts that were significantly activated by the genotoxic agent etoposide. Intriguingly, the differentially genes related to inflammatory and immune responses were enriched in resistant Capan-2 cells (**Figure 3-figure supplement 1A**), raising the possibility that pro-inflammatory factors may play a causal role in conferring chemoresistance. Gene set enrichment analysis further demonstrated a significant upregulation of genes associated with interferon-alpha (IFN-α) response, inflammatory response, and interferon-gamma (IFN-γ) response in etoposide-resistant cells (**Figure 3, A-C**). Along with BCL6 upregulation, the expression of IFN signaling-related genes was significantly increased accordingly (**Figure 3, D-E**).

**Figure 3.**
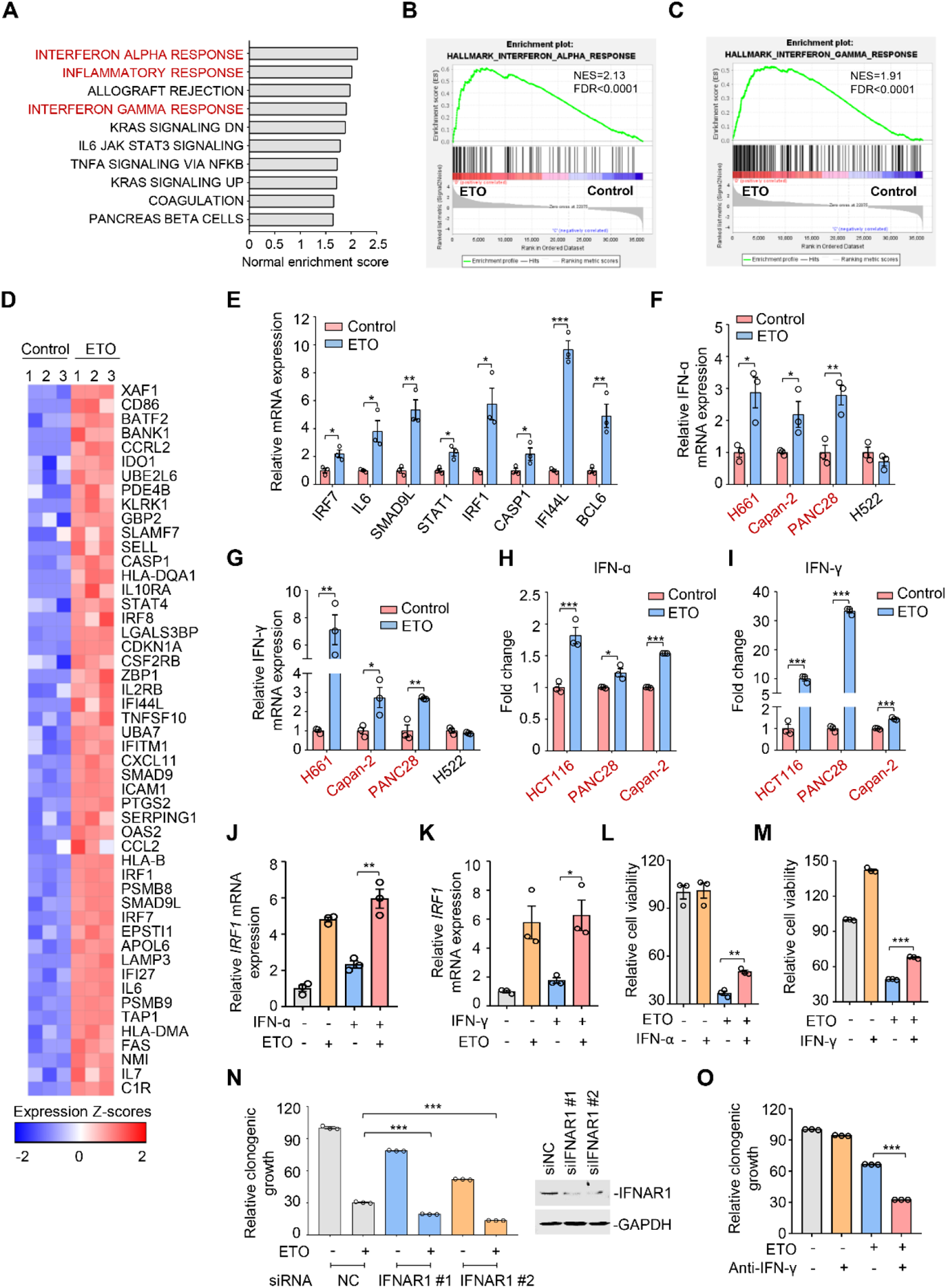
Genotoxic stress activates interferon responses. **(A)** Gene set enrichment analysis of pathways significantly upregulated in Capan-2 cells treated with 50 μM etoposide for 24 h (*n* = 3). GO analysis are shown in **Figure 3-figure supplement 1A.** (**B** and **C**) Enrichment plots for genes associated with interferon α (IFN-ɑ, **B**) and interferon γ (IFN-γ, **C**) responses in etoposide-treated Capan-2 cells. (**D**) Heap map illustrating of representative gene expression of IFN-ɑ and IFN-γ responses in treated Capan-2 cells. Z-scores were calculated based on counts of exon model per million mapped reads. Upregulated and downregulated genes were identified by a cutoff of *P* < 0.05. (**E**) Validation of expression of inflammation-related genes in (**D**). Capan-2 cells were treated with 50 μM etoposide for 24 h. QPCR assays were subsequently performed. Values are expressed as mean ± SEM of three independent experiments. \**P* < 0.05, \*\**P* < 0.01 and \*\*\**P* < 0.001, unpaired, two tailed *t*-test. (**F** and **G**) IFN-ɑ (**F**) and IFN-γ (**G**) mRNA expression levels in treated cells. ETO-sensitive and -resistant cells were treated with etoposide at their respective 1/2 IC_50_s for 24 h, and qPCR analysis was further performed. Values are expressed as mean ± SEM of three independent experiments. \**P* < 0.05, \*\**P* < 0.01, unpaired, two tailed *t*-test. ETO-resistant cell lines are marked in red. (**H** and **I**) IFN-ɑ (**H**) and IFN-γ (**I**) production in ETO-resistant cells. Cells were treated with etoposide at their respective 1/2 IC_50_s for 48 h. The concentrations of IFN-ɑ and IFN-γ in cell lysates were measured using an ELISA assay. Values are expressed as mean ± SEM of three independent experiments. \**P* < 0.05, \*\*\**P* < 0.001, unpaired, two tailed *t*-test. (**J** and **K**) Relative IRF1 mRNA levels in Capan-2 cells. Capan-2 cells were treated with 50 ng/mL IFN-ɑ (**J**) or 10 ng/mL IFN-γ (**K**) in the presence or absence of 50 μM etoposide. IRF1 mRNA levels were detected by qPCR assays. Values are expressed as mean ± SEM of three independent experiments. **\****P* < 0.05, \*\**P* < 0.01, unpaired, two tailed *t*-test. (**L** and **M**) Relative cell viability. ETO-sensitive H522 cells were treated with etoposide alone, 50 ng/mL IFN-ɑ (**L**), 10 ng/mL IFN-γ (**M**) or their combinations. Cell viability were examined by SRB assays. Values are expressed as mean ± SEM of three independent experiments by setting the control group as 100%. \*\**P* < 0.01, \*\*\**P* < 0.001, unpaired, two tailed *t*-test. (**N**) Clonogenic growth of Capan-2 cells treated with siIFNAR1, 0.4 μM etoposide, or their combinations (*left*). IFNAR1 silencing efficiency was examined using immunoblotting analysis (*right*). Data are expressed as mean ± SEM of three independent experiments. \*\*\**P* < 0.001, unpaired, two tailed *t*-test. Cell viability curves are shown in **Figure 3-figure supplement 1B.** (**O**) Clonogenic growth showing the relative survival of PANC28 cells treated with 0.2 μM etoposide, 10 μg/mL anti-IFN-γ or both. Values are expressed as mean ± SEM of three independent experiments. \*\*\**P* < 0.001, unpaired, two tailed *t*-test. Cell viability curves are shown in **Figure 3-figure supplement 1C**. The following source data and figure supplements are available for figure 3: **Figure 3-Source data 1.** Genotoxic stress activates interferon responses. **Figure 3-figure supplement 1.** Genotoxic stress activates interferon responses.

Recent work has revealed that consistent DNA damage triggers an inflammatory cytokine secretory phenotype in cultured cells (Rodier et al., 2009). To corroborate whether IFN-α and IFN-γ were similarly induced because of genotoxic agents, we assayed the gene expression of IFN-α and IFN-γ in treated cells. Our results showed that etoposide exposure resulted in an evident upregulation of IFN-α (**Figure 3F**) and IFN-γ transcription (**Figure 3G**) in etoposide-resistant H661, Capan-2, and PANC28 cells, but not in etoposide-sensitive H522 cells. We further examined the cellular production of IFN-α and IFN-γ in treated cells using a direct enzyme-linked immunosorbent assay, and found that etoposide treatment evoked a significant increase in IFN-α and IFN-γ contents in resistant cells (**Figure 3, H-I**).

Interferon regulatory factor 1 (IRF1), a key transcription factor that regulates cell proliferation and immune responses, is an inducible gene of type I and type II interferon (Castellaneta et al., 2014; Dery et al., 2014). To explore the effects of etoposide on IFN signaling, we examined IRF1 expression in resistant cells. We found that etoposide not only triggered a notable increase in IRF1 transcription itself, but also dramatically enhanced IFN-α- and IFN-γ-induced IRF1 expression in resistant cells (**Figure 3, J-K**), indicating the potent effect of etoposide on cellular interferon responses.

We next investigated the biological significance of IFN upregulation in the process of tumor adaptive response to genotoxic agents. Our results showed that exogenous addition of IFN-α and IFN-γ protected sensitive cells from etoposide-induced cell death (**Figure 3, L-M**). In contrast, siRNA knockdown of the IFN-α receptor IFNAR1 led to an enhanced sensitivity of resistant cells to etoposide, as indicated by impaired clonogenic growth (**Figure 3N**) and decreased etoposide IC_50_ values (**Figure 3-figure supplement 1B**). In line with these observations, antibodies against IFN-γ increased the killing ability of etoposide towards resistant cells (**Figure 3O** and **Figure 3-figure supplement 1C**). These results indicate that IFN activation provoked by genotoxic stress promotes tumor cell survival, leading to a tumor resistance phenotype.

### The interferon/STAT1 axis directly regulates BCL6 expression

Accumulating evidences show that IFNs produce pro-survival effects and mediate non-immune resistance to chemotherapy primarily through the transcriptional factor STAT1 (Khodarev et al., 2007; Minn, 2015). Following this direction, we examined STAT1 expression in treated cells and found that etoposide treatment promoted STAT1 protein abundance in etoposide-resistant PANC28 and HCT116 cells, but not in sensitive H522 cells (**Figure 4A**). Furthermore, genetic knockdown of STAT1 synergized with genotoxic agents to inhibit the clonogenic growth of resistant cells (**Figure 4B**). These results collectively suggest that the interferon/STAT1 axis is required for the therapeutic efficacy of etoposide and plays an essential role in tumor response to genotoxic stress.

**Figure 4.**
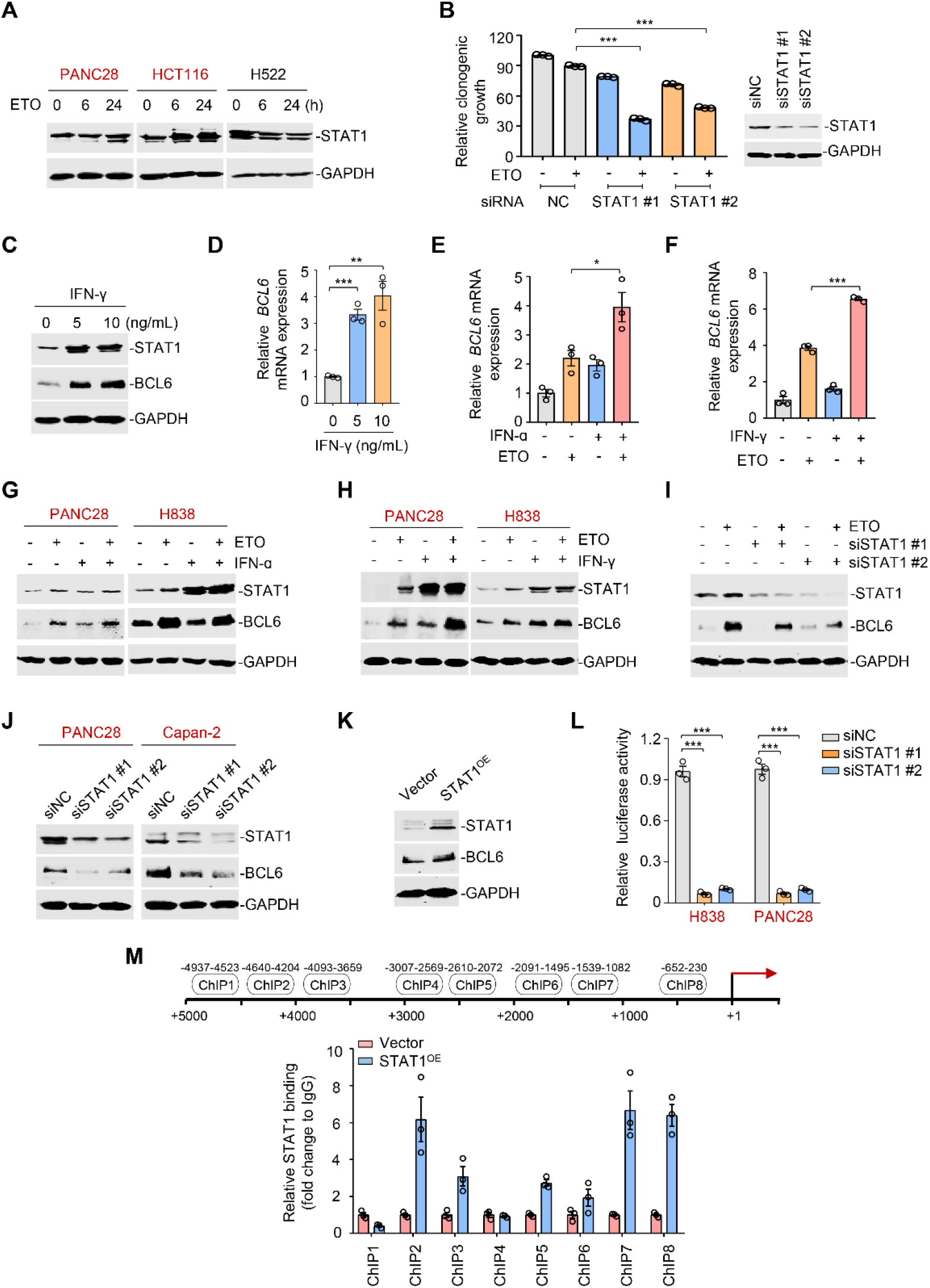
The interferon/STAT1 axis directly regulates BCL6 expression. (**A**) STAT1 protein levels by immunoblotting analysis. ETO-resistant and -sensitive cells were treated with etoposide at their respective 1/2 IC_50_s for indicated time points. Cell lysates were collected and subjected to immunoblotting analysis. ETO-resistant cell lines are marked in red. (**B**) Clonogenic growth of H838 cells treated with siRNAs targeting STAT1, 0.2 μM etoposide, or their combinations. STAT1 silencing efficiency was examined using immunoblotting analysis (*right*). Values are expressed as mean ± SEM of three independent experiments. \*\*\**P* < 0.001, unpaired, two tailed *t*-test. (**C**) IFN-γ increased BCL6 and STAT1 protein levels. H838 cells were treated with 5 or 10 ng/mL IFN-γ for 24 h. Cell lysates were subjected to immunoblot analysis with indicated antibodies. (**D**) Relative BCL6 mRNA expression. H838 cells were treated with 5 or 10 ng/mL IFN-γ for 24 h. BCL6 mRNA levels were detected by qPCR assays. Values are expressed as mean ± SEM of three independent experiments. \*\**P* < 0.01, \*\*\**P* < 0.001, unpaired, two tailed *t*-test. (**E** and **F**) Relative BCL6 mRNA expression. H838 cells were treated with 50 ng/mL IFN-ɑ (**E**) or 10 ng/mL IFN-γ (**F**) in the presence or absence of 50 μM etoposide. BCL6 mRNA levels were detected. Values are expressed as mean ± SEM of three independent experiments. \**P* < 0.05, \*\*\**P* < 0.001, unpaired, two tailed *t*-test. The same experiments were also repeated in Capan-2 cells (see **Figure 4-figure supplement 1A-B**). (**G** and **H**) Immunoblotting analysis for BCL6 and STAT1 protein expression. PANC28 or H838 cells were treated with 50 ng/mL IFN-ɑ (**G**) or 10 ng/mL IFN-γ (**H**) in the presence or absence of etoposide for 48 h. Cell lysates were subjected to immunoblotting analysis with specific antibodies against BCL6, STAT1 and GAPDH. (**I**) STAT1 knockdown impaired etoposide-induced BCL6 activation. STAT1 silencing was performed by RNA interference in H838 cells. Transfected cells were treated with 50 μM etoposide for 24 h, and cell lysates were subjected to immunoblotting analysis. (**J**) Silencing of STAT1 decreased BCL6 expression in ETO-resistant PANC28 and Capan-2 cells. (**K**) Overexpression of STAT1 increased BCL6 expression in Capan-2 cells. (**L**) Relative luciferase activity. siRNAs targeting STAT1 and BCL6n-luc vector were transiently co-transfected into ETO-resistant H838 and PANC28 cells. Luciferase activity was measured 48 h post-transfection. Bar graphs represent the mean ± SEM of three independent experiments. \*\*\**P* < 0.001, unpaired, two tailed *t*-test. (**M**) ChIP-qPCR data showing the enrichment of STAT1 binding to the BCL6 promoter region in Capan-2 cells. Values are expressed as mean ± SEM of three independent experiments. The experiments were performed twice. The following source data and figure supplements are available for figure 4: **Figure 4-Source data 1.** The interferon/STAT1 axis directly regulates BCL6 expression. **Figure 4-figure supplement 1.** The interferon/STAT1 axis directly regulates BCL6 expression.

Activated STAT1 drives an interferon-related gene signature for DNA damage tolerance (Minn, 2015), which prompted us to hypothesize that the interferon/STAT1 axis might regulate BCL6 expression. Given that IFN-γ activated IFN-stimulated gene expression specifically through the classical Janus kinase/STAT1 signaling, we first incubated resistant cells with 5 or 10 ng/mL recombinant IFN-γ, and found that IFN-γ significantly evoked a simultaneous increase in STAT1 and BCL6 protein expression (**Figure 4C**), implying that these two factors might be functionally linked. When noted, IFN-γ at the same concentrations evidently triggered BCL6 mRNA expression (**Figure 4D**). To further clarify the role of interferon signaling in modulating BCL6 expression, we treated resistant cells with etoposide in combination with IFN-α or IFN-γ, respectively. Our results showed that etoposide-mediated BCL6 transactivation could be further enhanced in etoposide-resistant H838 cells (**Figure 4, E-F**) and Capan-2 cells (**Figure 4-figure supplement 1A-B**) by the addition of IFN-α or IFN-γ. Moreover, etoposide induced STAT1 and BCL6 protein expression in resistant cells, whereas these effects could be potentiated by IFN-α (**Figure 4G**) or IFN-γ addition (**Figure 4H**), implying that etoposide-induced type 1 and type 2 interferon responses are required for STAT1 and BCL6 activation. Importantly, an increased expression of BCL6 by etoposide was apparently suppressed by STAT1 genetic silencing (**Figure 4I**). These results collectively suggest that etoposide transactivates BCL6 primarily through the interferon/STAT1 signaling pathway.

To elucidate the regulatory link of STAT1 on BCL6, we silenced STAT1 and found that STAT1 knockdown led to a marked decrease in BCL6 protein expression (**Figure 4J**), while STAT1 overexpression apparently increased BCL6 protein abundance (**Figure 4K**), implying that STAT1 may be upstream of BCL6. To elucidate whether STAT1 is a direct regulator of BCL6, we constructed a whole BCL6 promoter luciferase reporter and found that STAT1 interference resulted in a decreased BCL6 reporter activity (**Figure 4L**). Our chromatin immunoprecipitation coupled with qPCR analysis further revealed the recruitment of STAT1 to three putative binding regions of the BCL6 locus (**Figure 4M**). These results reinforced the direct regulation of the interferon/STAT1 signaling pathway on BCL6 expression.

### The tumor suppressor PTEN is a functional target of BCL6

After characterizing STAT1 as an upstream regulator of BCL6, we next explored BCL6 downstream signaling responsible for adaptive response to genotoxic stress. Considering two lines of evidences showing that: (1) phosphatase and tensin homolog (PTEN), a lipid phosphatase that antagonizes the phosphatidylinositol 3-kinase pathway (Lee, Chen, & Pandolfi, 2018), was enriched in BCL6 promoter binding peaks in primary germinal center B cells (Ci et al., 2009), and that (2) BCL6 directly binds to the promoter locus of PTEN in patient-derived acute lymphoblastic leukemia (Geng et al., 2015), we hypothesized that an increase in BCL6 expression by genotoxic stress might inhibit PTEN and subsequently promote cell survival. To this end, we performed transcriptome analysis and found an evident decrease in PTEN expression in Capan-2 and H661 cells exposed to etoposide (**Figure 5A**). The analysis of datasets from TCGA further revealed that PTEN deletion was mutually exclusive with BCL6 amplification (**Figure 5B**). Furthermore, our qPCR (**Figure 5C**) and immunoblotting analysis (**Figure 5D**) showed that the upregulation of BCL6 was accompanied by a decreased expression of PTEN at both the mRNA and protein levels. To further support the notion that BCL6 repressed the expression of PTEN, we overexpressed BCL6 and observed a significant decrease in PTEN (**Figure 5E**). In contrast, doxycycline-inducible knockdown of BCL6 increased PTEN expression (**Figure 5F**). Our ChIP-qPCR data further revealed that etoposide treatment significantly increased the occupancy of BCL6 at the promoter region of *PTEN* (**Figure 5G**). These results indicated that PTEN is a functional target of BCL6 and largely contributes to genotoxic stress tolerance in tumor cells.

**Figure 5.**
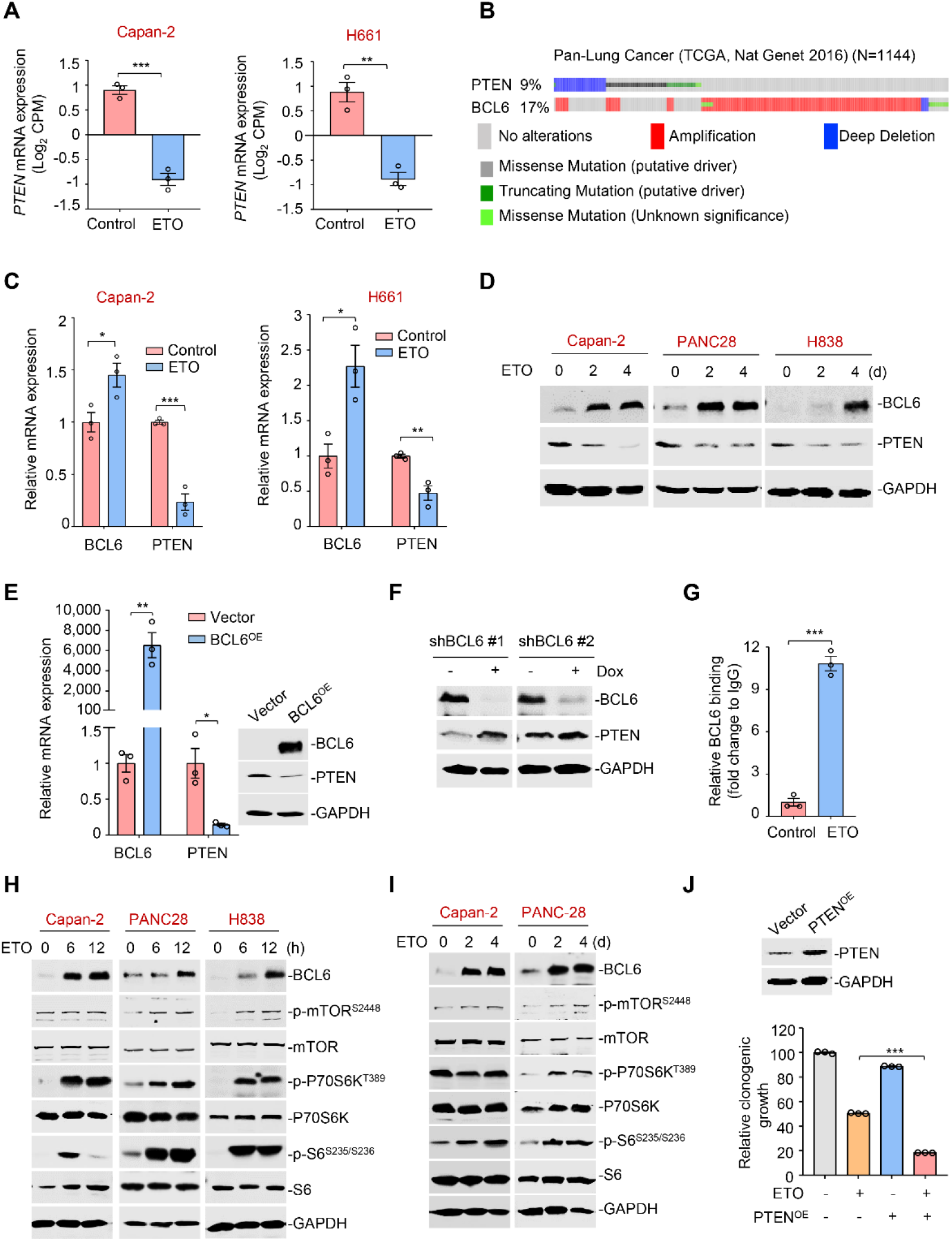
The tumor suppressor PTEN is a functional target of BCL6. (**A**) Normalized PTEN expression levels in etoposide-resistant Capan-2 and H661 cells treated with etoposide at their respective IC_50_s for 24 h. RNA-seq tag count at exons was plotted as counts of exon model per million mapped reads. Values are expressed as mean ± SEM of three independent experiments. \*\**P* < 0.01, \*\*\**P* < 0.001, unpaired, two tailed *t*-test. (**B**) Genomic alteration of BCL6 and PTEN according to TCGA database (*n* = 1144). The percentage of gene alteration is shown. (**C**) Relative mRNA expression of BCL6 and PTEN. Capan-2 and H661 cells were exposed to etoposide at their respective 1/2 IC_50_s for 24 h. QPCR analysis was further carried out. Values are expressed as mean ± SEM of three independent experiments. \**P* < 0.05, \*\**P* < 0.01, \*\*\**P* < 0.001, unpaired, two tailed *t*-test. (**D**) BCL6 and PTEN protein levels in Capan-2, PANC-28 and H838 cells. Cells were treated with etoposide at their respective 1/4 IC_50_s for 2 or 4 days. Cell lysates are subjected to immunoblotting analysis. (**E**) BCL6 overexpression decreased PTEN mRNA and protein levels in HCT116 cells. Cells were transfected with pcDNA3.1-BCL6 or pcDNA3.1 control plasmid. Total mRNA and protein were extracted and subjected to qPCR analysis (*left*) and immunoblotting analysis (*right*). Values are expressed as mean ± SEM of three independent experiments. \**P* < 0.05, \*\**P* < 0.01, unpaired, two tailed *t*-test. (**F**) BCL6 inducible knockdown increased PTEN expression. Immunoblotting analysis of PTEN in HCT116 cells treated with or without doxycycline. (**G**) BCL6 binding levels at the promoter region of *PTEN* examined by ChIP-qPCR assays. Values are expressed as mean ± SEM of three independent experiments. \*\*\**P* < 0.001, unpaired, two tailed *t*-test. (**H**) Etoposide activated mTOR signaling components in etoposide-resistant Capan-2, PANC-28 and H838 cells. Cells were treated with etoposide at their respective 1/2 IC_50_s for 6 or 12 h. Whole-cell lysates were prepared and subjected to immunoblotting analysis. (**I**) A long-term treatment with etoposide activated mTOR signaling components in ETO-resistant cells. Capan-2 and PANC-28 cells were treated with 10 μM etoposide for 2 or 4 days. Cell lysates were subjected to immunoblotting analysis. (**J**) PTEN overexpression increased the sensitivity of etoposide-resistant cells to etoposide. PANC28 cells were transfected with pCDH-PTEN or pCDH control plasmid. PTEN overexpression efficiency was measured immunoblotting analysis (*up*). Quantification of clonogenic growth after 7 days treatment with 0.2 μM etoposide (*down*). Values are expressed as mean ± SEM of three independent experiments. \*\*\**P* < 0.001, unpaired, two tailed *t*-test. The following source data are available for figure 5: **Figure 5-Source data 1.** The tumor suppressor PTEN is a functional target of BCL6.

It is well-known that PTEN acts as a tumor suppressor and hampers the activation of the proto-oncogenic mTOR pathway (Martelli et al., 2011). We further explored the effects of etoposide treatment on the mTOR signaling. Our immunoblotting results showed that phosphorylation of mTOR (S2448), S6K (T389) and S6 (S235/S236) was strikingly increased, along with an aberrant BCL6 expression in etoposide-treated resistant cells (**Figure 5H**). Similar results were also obtained in a long-term drug exposure assay (**Figure 5I**). Notably, overexpression of PTEN enhanced the antitumor effects of etoposide (**Figure 5J**). These results collectively suggest that the PTEN/mTOR pathway is a downstream signaling of BCL6.

### BCL6 inhibition conquers resistance of cancer cells to genotoxic stress *in vitro*

Since tumor adaptive resistance to genotoxic stress was attributed to BCL6 transactivation, we tested whether pharmacological inhibition of BCL6 could restore the sensitivity of resistant cancer cells to genotoxic agents. We suppressed BCL6’s function using two BCL6 pharmacological inhibitors, BI3802 and compound 7. BI3802 was reported as a BCL6 degrader (Kerres et al., 2017; Slabicki et al., 2020), while compound 7 targeted the BCL6 BTB/POZ domain and prevented its partner binding (Kamada et al., 2017). Our results showed that multiple resistant cell lines became vulnerable to etoposide in the presence of BI3802 or compound 7 (**Figure 6A**). In addition, BI3802 addition could shift the IC_50_ values of doxorubicin (**Figure 6-figure supplement 1A**). Moreover, combination index values (CIs) were further employed to indicate drug synergy, and our results showed that the majority of CIs at 50%, 75% and 90% of the effective dose of each drug pair (etoposide plus BI3802, or etoposide plus compound 7) in five resistant cell lines were all lower than 1 (**Figure 6B**), displaying a synergistic action of etoposide and BCL6-targeted therapy. We further assessed the combined effects of etoposide and the BCL6 inhibitor BI3802 in a long-term colony-formation assay. Our results showed that the combination of etoposide and BI3802 led to a robust growth inhibition of cultured colonies (**Figure 6C**). As expected, addition of BI3802 markedly enhanced the inhibitory effects of etoposide on soft-agar colony formation (**Figure 6D**). A combinative synergy was also obtained for doxorubicin and targeted BCL6 inhibition (**Figure 6-figure supplement 1B-C**). All these data indicate that BCL6 blockage could restore the sensitivity of cancer cells to genotoxic agents.

**Figure 6.**
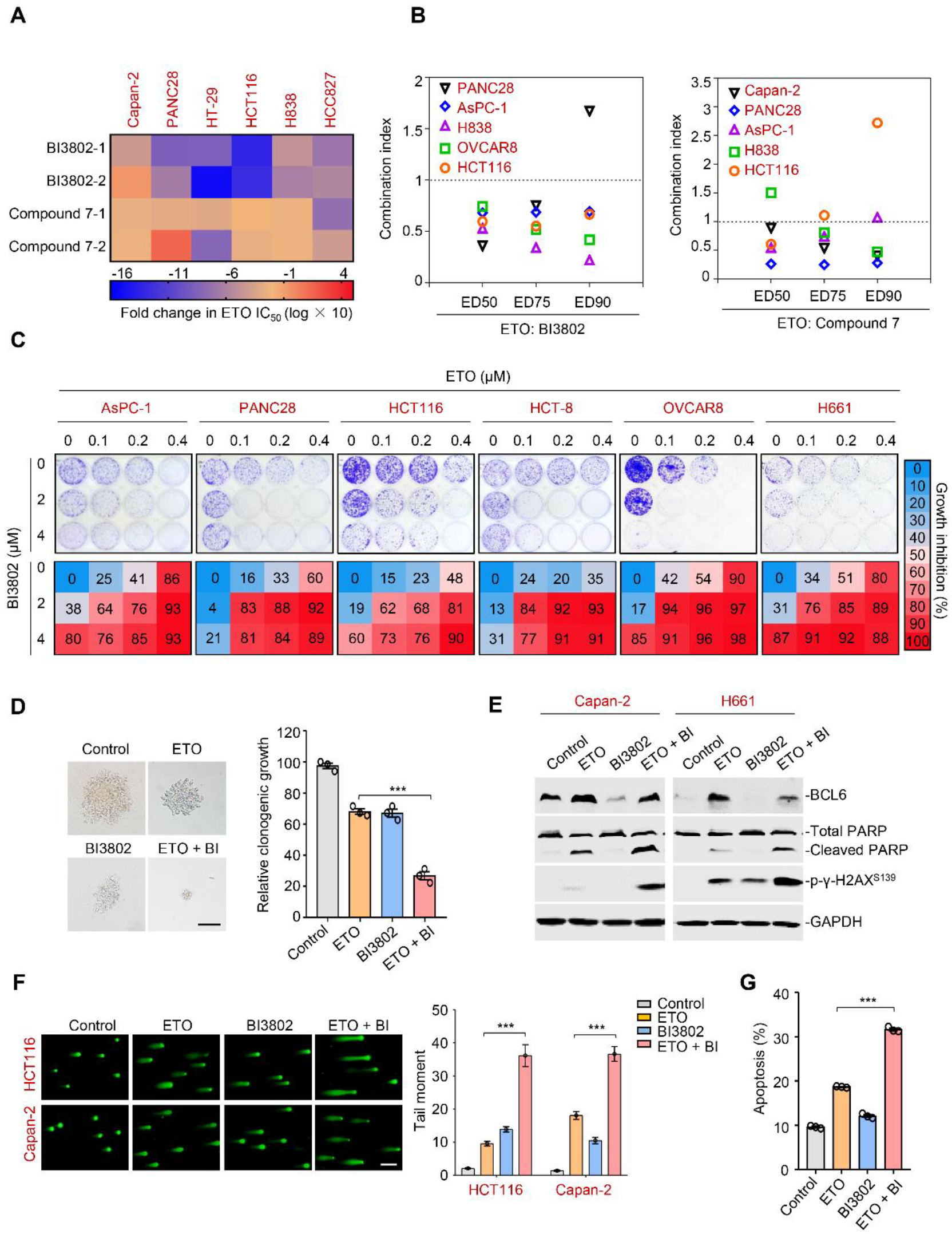
Therapeutic suppression of BCL6 sensitizes genotoxic agents. (**A**) Pharmacological inhibition of BCL6 increased ETO sensitivity. Various types of cancer cells were treated with etoposide at gradient concentrations for 48 h in the presence of 10 μM BI3802 or 20 μM Compound 7 (*n* = 2 biological replicates). IC_50_s were measured using SRB assays. For graphs, log(IC_50_) of control cells was subtracted from log(IC_50_) of BI3802 or Compound 7-treated cells and multiplied by ten to be depicted as log fold change ×10. Targeted inhibition of BCL6 also increased ADR sensitivity (see **Figure 6-figure supplement 1A**). (**B**) Synergistic interaction between BCL6 inhibitors (BI3802 or Compound 7) and ETO. Growth inhibition was averaged and input into CalcuSyn software to extrapolate combinational index values (CI) at 50% effective dose (ED50), 75% effective dose (ED75) and 90% effective dose (ED90). CI values < 1 represent synergism. The synergy between BI3802 and ADR was also detected in H838, Capan-2 and AsPC-1 cells (see **Figure 6-figure supplement 1B**). (**C**) Inhibition of clonogenic growth by the combined regimen. Representative long-term clonogenic images (*up*) and quantified clonogenic growth inhibition results (*down*) for cells treated with ETO, BI3802, or their combinations. Data are presented as mean of three independent experiments. The same experiments were also conducted for ADR (see **Figure 6-figure supplement 1C**). (**D**) Inhibition of soft-agar colony growth by the combined regimen. HCT116 cells were exposed to 0.2 μM etoposide, 2 μM BI3802, or their combinations. Representative images of soft-agar colonies (*left*) and the relative clonogenic growth (*right*) are shown. Scale bar, 100 μm. Values are expressed as the mean ± SEM of three independent experiments. \*\*\**P* < 0.001, unpaired, two tailed *t*-test. (**E**) Immunoblotting analysis showing the protein expression of BCL6, p-γ-H2AX^S139^ and cleaved-PARP in Capan-2 and H661 cells treated with 15 μM etoposide, 10 μM BI3802 or their combinations for 48 h. Cell lysates were subjected to immunoblotting analysis. (**F**) Comet assays. HCT116 and Capan-2 cells were treated with etoposide, BI3802, or their combinations for 48 h. The tail moment was quantified for 50 cells for each experimental condition (*right*). Scale bar, 100 μm. Values are expressed as mean ± SEM. \*\*\**P* < 0.001, unpaired, two tailed *t*-test. (**G**) Quantification of apoptotic cells in Capan-2 cells analyzed by flow cytometry. Cells were exposed to 15 μM etoposide, 10 μM BI3802 or their combinations for 48 h. Percentage of positive cells was counted. Values are expressed as mean ± SEM of three independent experiments. \*\*\**P* < 0.001, unpaired, two tailed *t*-test. The following source data and figure supplements are available for figure 6: **Figure 6-Source data 1.** Therapeutic suppression of BCL6 sensitizes genotoxic agents. **Figure 6-figure supplement 1.** BCL6 inhibition sensitizes cancer cells to doxorubicin.

DNA damage potency triggered by genotoxic agents is a determinant of tumor response to chemotherapy (Bouwman & Jonkers, 2012; Pearl, Schierz, Ward, Al-Lazikani, & Pearl, 2015). The accumulation of DNA damage further causes genome instability and consequently triggers cell apoptosis (Broustas & Lieberman, 2014). The fact that BCL6 upregulation was associated with reduced phosphorylated H2AX levels in HCT116 xenografts (**Figure 2E**) prompted us to explore whether the targeted inhibition of BCL6 could promote DNA damage in the presence of genotoxic agents. Our results showed that the combined regimen of etoposide and BI3802 resulted in more poly (ADP-ribose) polymerase cleavage and a higher phosphorylated H2AX expression (Ser139) than single agent alone (**Figure 6E**). In addition, more DNA damage occurred as indicated by a significantly higher tail moment observed in a comet assay in the combined treatment group (**Figure 6F**). Consequently, an increase in the number of apoptotic cells was observed in the drug pair group (**Figure 6G**). Taken together, these data suggest that BCL6 blockade potentiates genotoxic agents by inducing DNA damage and growth inhibition.

### Targeted inhibition of BCL6 sensitizes genotoxic agents *in vivo*

We next investigated whether our combined therapeutic approach is effective in tumor preclinical mouse models. BI3802 was reported to possess a poor bioavailability (Kerres et al., 2017). Therefore, we applied FX1, another BCL6 pharmacological inhibitor, which disrupts the interaction between BCL6 and co-repressors with satisfactory antitumor effects *in vivo* (Beguelin et al., 2016; Cardenas et al., 2016). We first set up a xenograft mouse model using HCT116 cells. Once the average volume of xenografts reached ∼100 mm^3^, mice were treated with etoposide or the vehicle. We found that BCL6 was upregulated at both mRNA and protein levels in xenografts as early as 2 days after drug administration, and intriguingly, this effect was sustained during the treatment period (**Figure 7A**), implying the emergence of a resistance phenotype. Strikingly, the addition of FX1 from day 2 significantly enhanced the therapeutic potency of etoposide, as indicated by a decreased tumor volume and tumor weight (**Figure 7B**). Most importantly, administration of 10 mg/kg etoposide and 5 mg/kg FX1 was well-tolerated in mice since the levels of blood biochemical indicators were marginally affected (**Supplementary Table 4**).

**Figure 7.**
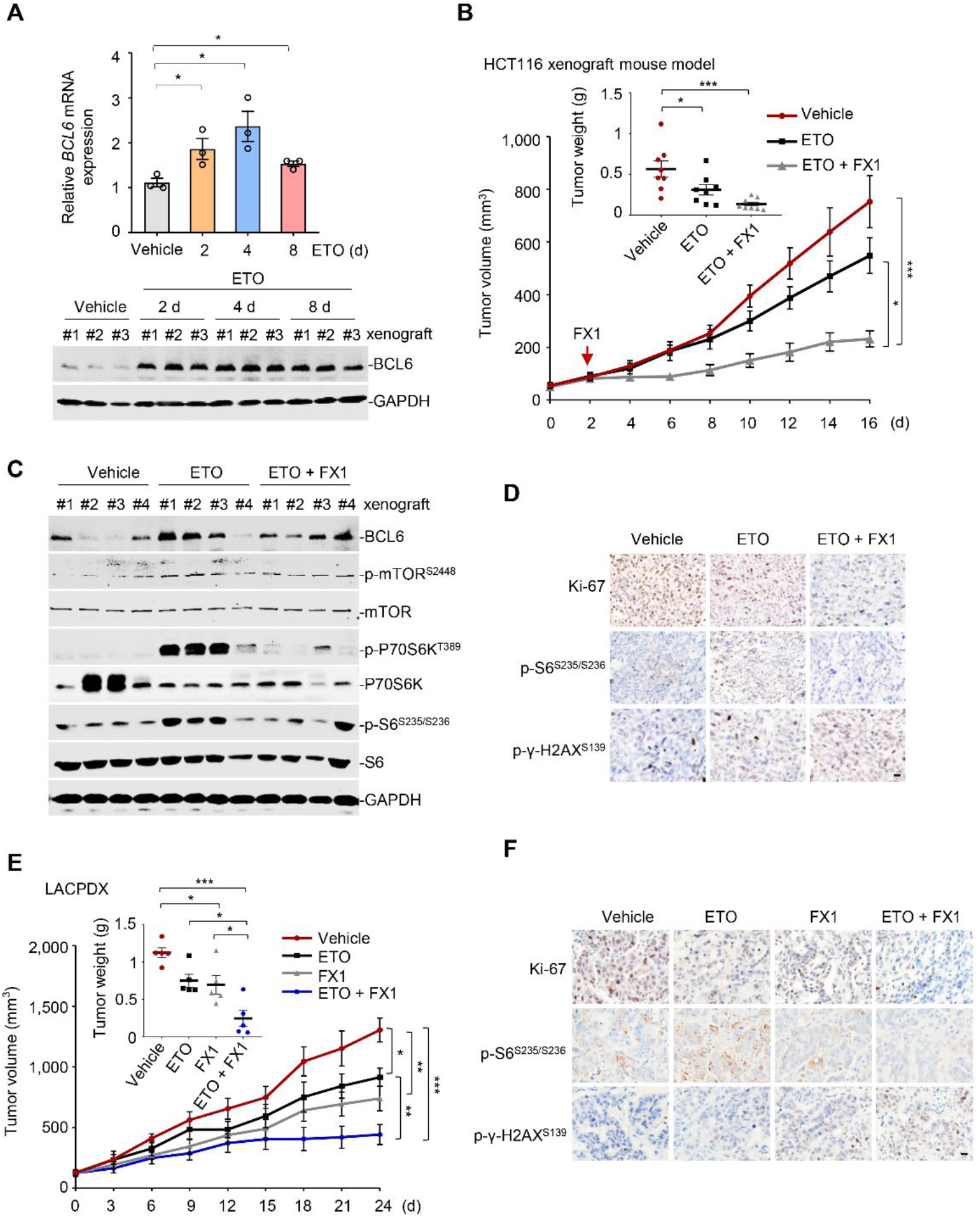
Pharmacological inhibition of BCL6 synergizes etoposide *in vivo*. (**A**) Etoposide increased BCL6 mRNA (*up*) and protein (*down*) expression in HCT116 xenografts. Tumor tissues were isolated on day 2, 4 or 8 after etoposide treatment. QPCR and immunoblotting analysis for BCL6 expression were conducted. BCL6 mRNA expression values represent mean of three independent replicates ± SEM. \**P* < 0.05, unpaired, two tailed *t*-test. (**B**) Tumor growth curves. Mice bearing HCT116 xenografts were treated with vehicle, etoposide (10 mg/kg body weight), and etoposide plus FX1 (5 mg/kg body weight) for indicated times. Average tumor weight on day 16 is shown in the inset. Values are expressed as mean ± SEM, *n* = 8. \**P* < 0.05, \*\*\**P* < 0.001, one-way ANOVA with Tukey’s multiple-comparisons test. (**C**) Protein expression of BCL6 and mTOR signaling components in HCT116 xenografts. Tumors were harvested at the end of treatment and subjected to immunoblotting analysis. Four biologically independent samples per group are shown. (**D**) Representative immunohistochemical staining of tumors in HCT116 xenografts. Tumor tissues from HCT116 xenografts on day 16 were examined for the expression of Ki-67, p-γ-H2AX^S139^, and p-S6^S235/S236^. Scale bar, 50 µm. (**E**) Tumor growth curves. Mice bearing primary *KRAS*-mutant lung cancer xenografts (LACPDX) were treated with vehicle, etoposide (10 mg/kg body weight), FX1 (5 mg/kg body weight) or both drugs for 24 days. Average tumor weight on day 24 is shown in the inset (*n* = 5). Values are expressed as mean ± SEM. \**P* < 0.05, \*\**P* < 0.01, \*\*\**P* < 0.001, one-way ANOVA with Tukey’s multiple-comparisons test. (**F**) Representative immunohistochemical staining of LACPDX tumors. Tumor tissues from LACPDX on day 24 were evaluated for the expression of Ki-67, p-S6^S235/S236^ and p-γ-H2AX^S139^. Scale bar, 50 µm.

Immunoblot analysis of tumor lysates revealed a marked increase in p-mTOR (S2448), p-P70S6K (T389), and p-S6 (S235/S236) expression levels in etoposide-treated xenografts (**Figure 7C**), whereas addition of FX1 suppressed the activation of the mTOR signaling pathway. Immunohistochemistry analysis additionally showed an increase in p-H2AX (S139) expression and weaker Ki-67 signals in the xenografts from the drug pair group (**Figure 7D**), suggesting a fundamental role of BCL6-targeted therapy in sensitizing etoposide *in vivo*.

To evaluate the antitumor activity of FX1+ etoposide in a more clinically relevant mouse model, we established a patient-derived xenograft model of lung adenocarcinoma harboring a G12V mutation in KRAS (LACPDX). Our results showed that the combination of etoposide and FX1 significantly suppressed tumor weight and tumor volume compared with single agent alone (**Figure 7E**), without causing systemic toxicity in mice (**Supplementary Table 5**). In agreement, addition of FX1 markedly decreased p-S6 (S235/S236) expression and increased p-H2AX (S139) expression in LACPDX (**Figure 7F**). These results collectively suggest that BCL6 is a crucial combinatorial target in the sensitization of resistant tumors to genotoxic agents *in vivo*.

## Discussion

The exploration of underlying resistance mechanisms of genotoxic agents may allow the prediction of patient responses, the design of rational combination therapies and the implementation of re-sensitization strategies. Here, we show that BCL6 upregulation is a prominent mechanism to protect tumor cells from genotoxic killing. Our current findings support the notion that BCL6 functions as a central factor in mediating therapy resistance: (1) the interferon/STAT1 pathway serves as an upstream regulator of BCL6, (2) the tumor suppressor PTEN is identified as a functional target of BCL6, (3) the activation of BCL6 signaling leads to a sustained pro-survival phenotype, whereas blocking it enhances the therapeutic efficacy of genotoxic agents. Our findings further establish a rationale for the concurrent targeting of BCL6 to conquer tumor tolerance to genotoxic stress, as evidenced by the striking synergy of genotoxic therapy and BCL6-targeted therapy *in vitro* and *in vivo* (**Figure 6** and **Figure 7**).

BCL6 acts as a gatekeeper to protect germinal center B cells from undergoing somatic hypermutation and class-switch recombination against DNA damage (Duy et al., 2010; Polo, Ci, Licht, & Melnick, 2008). In this study, we showed, for the first time, that BCL6 was markedly upregulated by genotoxic agents in both *in vitro* and *in vivo* settings, leading to a resistance phenotype (**Figure 1** and **Figure 2**). Furthermore, high BCL6 levels were positively associated with unfavorable clinical outcomes (**Figure 1G**). Our results were conceptually in line with recent findings showing that BCL6 enabled heat stress tolerance in vertebrates (Fernando et al., 2019) and conferred tyrosine kinase inhibitor resistance in Ph^+^ acute lymphoblastic leukemia (Duy et al., 2011). As reported in our recent work (Guo et al., 2021), BCL6 activation attenuated the antitumor efficacy of clinical BET inhibitors in *KRAS*-mutant lung cancers. Combining these findings together, we speculate that BCL6 may functionally program tumor pro-survival signals in drug response and can be used as a predictive biomarker for therapy resistance. As an essential transcription repressor, BCL6 suppresses rapid proliferation and survival of germinal center B cells by recruiting co-repressors, such as BCOR, NCOR and SMRT, to its BTB domain (Huang, Hatzi, & Melnick, 2013). Therapy targeting the BCL6 BTB domain lateral groove displayed inhibitory effects in the treatment of lymphoma (Cheng et al., 2018). Based on the substantial role of BCL6 in the tumor adaptive response to drug treatments, we assessed the therapeutic efficacy of BCL6-targeted therapy in combination with etoposide, which markedly strengthened DNA damage and tumor growth inhibition (**Figure 6** and **Figure 7**), without causing obvious toxicity in mice, providing a combinatorial strategy with translational significance.

BCL6 upregulation is required for maintaining B cells in germinal center compartments (Basso & Dalla-Favera, 2012). Once expressed in B cells, BCL6 is tightly controlled through an auto-regulatory circuit model, in which BCL6 negatively regulates its own transcription by binding to its gene promoter (Kikuchi et al., 2000; Pasqualucci et al., 2003). We recently reported that BRD3 maintained the auto-regulatory circuit of BCL6 by directly interacting with BCL6. Aberrant genomic or expressional changes of BCL6 have been detected in lymphomas and multiple solid tumors, including breast cancer, glioblastoma or ovarian cancer (Walker et al., 2015; Y. Q. Wang et al., 2015; Xu et al., 2017). Limited lines of evidence have revealed that the transcriptional factor STAT5 serves as a direct negative regulator of BCL6 in lymphomas (Walker, Nelson, & Frank, 2007), and FoxO3a promoted BCL6 expression in leukemia cells exposed to BCR-ABL inhibitors (Duy et al., 2011; Fernandez de Mattos et al., 2004). However, the transcriptional regulation pattern of BCL6 in solid tumors remains unexplored. Our findings demonstrated that the genotoxic agent etoposide activated the interferon/STAT1 signaling axis, which directly upregulated BCL6 by recruiting STAT1 to the binding regions of the BCL6 locus (**Figure 3** and **Figure 4**). The phenomenon that BCL6 could be transactivated by STAT1 was partially observed in imatinib-treated chronic myeloid leukemia cells (Madapura et al., 2017). These findings collectively suggest that BCL6 may be concisely and dynamically regulated by a unique mechanism in the specific tumor context.

While numerous cell-intrinsic processes are known to play critical roles in tumor response to genotoxic agents, increasing attention has been paid to multiple cell-extrinsic components of the tumor microenvironment that influence the malignant phenotype and disease progression. During DNA damage, the production of cellular mitogenic growth factors and proteases, such as HGF, EGF, and MMP, are programmed to facilitate tumor growth (Bavik et al., 2006; Coppe et al., 2008). In addition to these pro-survival molecules, the production of pro-inflammatory cytokines (e.g., IL6) provoked by chemotherapy, will promote anti-apoptotic signaling and intrinsic chemo-resistance (Gilbert & Hemann, 2010; Poth et al., 2010). In this study, we showed that, in response to genotoxic stress, etoposide-resistant cells rapidly increased the production of IFN-α and IFN-γ, and more importantly, the increase in IFNs was sufficient to protect cells from genotoxic killing (**Figure 3**). These findings support the essential role of IFNs in the tumor microenvironment of conferring drug resistance, along with the fact that the IFN-related DNA damage resistance signature acts as a predictive marker for chemotherapy (Post et al., 2018; Weichselbaum et al., 2008). Our results further delineate a mechanism by which increased production of IFN-α or IFN-γ facilitated cancer cells to evade genotoxic stress by activating the transcriptional factor STAT1 (**Figure 4**). Although genotoxic therapy-induced damage to the tumor microenvironment promotes treatment resistance through cell nonautonomous effects (Sun et al., 2012), whether targeting the biologically notable upregulation of IFNs in conjunction with conventional therapy could enhance the treatment response still requires additional experimentation.

The BCL6 transcriptional program for the direct silencing of multiple target genes has been elaborated in primary B cells and lymphoma (Ci et al., 2009). However, few target genes of BCL6 have been characterized in solid tumors. Our study identified PTEN, the most frequently mutated tumor suppressor (Lee et al., 2018), as a functional target of BCL6 in therapy resistance (**Figure 5**). We showed that the overexpression of BCL6 suppressed PTEN, while the knockdown of BCL6 increased the expression of PTEN (**Figure 5, E-F)**. Furthermore, the combination of BCL6 inhibitors and genotoxic agents resulted in a marked suppression of the PTEN downstream component mTOR *in vivo* (**Figure 7, C-D**), reinforcing that mTOR activation is an actionable mechanism that confers drug resistance (Tanaka et al., 2011). When acting as a transcriptional repressor, the BCL6 BTB domain recruits the co-repressors NCOR, SMRT, and BCOR (Ghetu et al., 2008). The mechanism by which BCL6 mediated the repression of PTEN and whether this action is dependent on the BCL6 BTB domain still requires further investigation.

## Methods

### Cell lines and culture

H1975 (RRID:CVCL_1511), PC9 (RRID:CVCL_B260), H661(RRID:CVCL_1577), H522 (RRID:CVCL_1567), HCC827 (RRID:CVCL_2063), H838 (RRID:CVCL_1594), DLD-1 (RRID:CVCL_0248), HT-29 (RRID:CVCL_0320), HCT-8 (RRID:CVCL_2478), HCT116 (RRID:CVCL_0291), LoVo (RRID:CVCL_0399), AsPC-1 (RRID:CVCL_0152), BxPC-3 (RRID:CVCL_0186), Capan-2 (RRID:CVCL_0026), PANC28 (RRID:CVCL_3917), ES-2 (RRID:CVCL_3509), OVCAR8 (RRID:CVCL_1629), OVCA420 (RRID:CVCL_3935), HEY (RRID:CVCL_2Z96) and HEYA8 (RRID:CVCL_8878) were purchased from the American Type Culture Collection (Manassas, VA, USA). PANC-1 (RRID:CVCL_0480) and MIA PaCa-2 (RRID:CVCL_HA89) were purchased from the Shanghai Cell Bank of the Chinese Academy of Sciences (Shanghai, China). All cell lines were maintained in the appropriate culture medium supplemented with 10% fetal bovine serum and 100 U/mL penicillin/streptomycin. Experiments were performed with cells under 15 passages. All cell lines were authenticated by STR analysis and routinely tested for *mycoplasma* by using the Mycoalert Detection Kit (Beyotime, Jiangsu, China). The culture medium of cell lines is listed in **Supplementary Table 1**.

### Plasmids and reagents

The inducible BCL6 shRNA vectors were generated based on a pLVX-TetOne-Puro vector (RRID: Addgene 124797) according to standard protocols. All constructs were verified by sequencing. shRNAs sequence targeting BCL6 are available in **Supplementary Table 2**. Recombinant human IFN-α1 (z02866) was purchased from Genscript (Nanjing, China). Recombinant IFN-γ (300-02) and anti-human IFN-γ antibody (506532) were purchased from PeproTech (Rocky Hill, USA). Etoposide (HY-13629, a topoisomerase II inhibitor), doxorubicin (HY-15142, a topoisomerase II inhibitor), cisplatin (HY-17394, a DNA synthesis inhibitor), carboplatin (HY-17393, a DNA synthesis inhibitor), taxol (HY-B0015, a microtubule association inhibitor) and gemcitabine (HY-17026, a DNA synthesis inhibitor) were purchased from MedChemExpress (Monmouth Junction, USA).

### Cell viability assay

Cell viability was assessed using the sulforhodamine B (SRB) assay. Cells (2,000 - 5,000 cells per well) were seeded onto 96-well plates in appropriate cell culture medium, allowed to attach overnight, and treated with the indicated drug concentrations. Approximately 48 h later, the cells were fixed in 50% trichloroacetic acid at 4°C for 1 h, stained with 0.4% SRB, and dissolved in a 10 mM Tris solution. The absorbance (optical density, OD) was read at a wavelength of 515 nm. The IC_50_ values were calculated using GraphPad Prism 8.0 (RRID:SCR_002798), and the CI values were evaluated using CalcuSyn software (Version 2; Biosoft).

### Two-dimensional clonogenic assay

Cells (1,000-2,000 cells per well) were seeded onto 12-well plates. After 24 h, cells were treated with the indicated drug for about 7 - 10 days. When grown into visible clones, the cells were washed with phosphate-buffered saline (PBS), fixed with 4% paraformaldehyde, stained with 0.2% crystal violet and photographed. Stained cells were then dissolved in 10% acetic acid. The absorbance of the stained solution was read at a wavelength of 595 nm in a 96-well plate. The relative viability was calculated by setting that of untreated cells as 100%.

### Soft-agar colony formation assay

The soft-agar colony formation assay was conducted to evaluate the inhibitory effects of different treatments on the anchorage-independent growth of tumor cells. The bottom layer of soft agar was prepared by mixing 2 × growth medium and 1.5% noble agar (BD Biosciences, San Jose, CA) at a 1:1 ratio and the mixture was poured into 6-well plates. Cells (1,000 - 2,000 cells per well) were suspended in the second soft agar layer that contained 0.5% low melting point agar mixed with growth media and spread over the bottom layer. After solidification, the growth medium was added into each well. After incubation for 5-7 days, cells were treated with various drugs for 10-15 days. When grew into visible clones, cells were imaged using a fluorescence microscope and counted to evaluate cell viability.

### Cell apoptosis assays

Cell apoptosis was quantified using flow cytometry (FACSCalibur, BD) as described previously (Elkabets et al., 2013). For cell apoptosis, the cells exposed to drugs for the indicated times were washed twice with PBS, re-suspended in 400-500 μL of 1× binding buffer (BD), and stained with 5 μL of Annexin V–FITC and 5 μL of propidium iodide (PI, Sigma; 5 μg/mL) for 15 min at room temperature in the dark. Cells were detected using flow cytometry (FACS Calibur, BD) and quantitative analysis was carried out using FlowJo software (RRID:SCR_008520).

### RNA interference

For siRNA transfection, the cells were plated at a confluence of approximately 40%-60% in basal culture medium and transfected with siRNA duplex using Lipofectamine TM 2000 reagent (ThermoFisher Scientific) according to the manufacturer’s instructions for 6 h. After that, the medium of the transfected cells were replaced by complete medium, and the cells were plated into wells and exposed to the drugs. Western blotting was applied to detect the interference efficiency of target genes.

### RNA isolation and RT-qPCR analysis

Total RNA from cell lines was isolated using TRIzol extraction (Invitrogen). cDNA was then prepared using the PrimeScript RT reagent kit (TaKaRa). QPCR reactions were performed according to the manufacturer’s instructions using SYBR® Premix Ex Taq kit (TaKaRa). All reactions were performed in triplicates. The CT difference values between the target gene and housekeeping gene (*GAPDH*) were calculated using the standard curve method. The relative gene expression was calculated. The sequences of primers used for qPCR are listed in **Supplementary Table 3.**

### ChIP analysis

ChIPs were performed using cross-linked chromatin from Capan-2 cells and either anti-BCL6 antibodies (1;1000, Cell Signaling Technology Cat# 14895, RRID:AB_2798638), anti-STAT1 antibodies (1;1000, Abclonal Cat# A12075, RRID: AB_2758978), or normal rabbit IgG (CST, 2729), using SimpleChIP Plus Enzymatic Chromatin Immunoprecipitation kit (agarose beads) (Cell Signaling Technology, 9004). The enriched DNA was quantified by qPCR analysis using the primers listed in **Supplementary Table 3.**

### Western blotting assay

The preparation of cell lysis was performed according to standard methods. Cells were treated with the respective concentrations of drug for indicated times. Afterward, the cells were washed slightly with ice-cold PBS, and then lysed with radio-immunoprecipitation assay (RIPA) buffer containing protease and phosphatase inhibitor cocktail (Calbiochem). The protein concentrations of cell lysate supernatants were assayed using a BCA protein assay kit (Thermo Scientific). Protein samples were resolved on 8–12% SDS–polyacrylamide gels and transferred to nitrocellulose membranes (Millipore). Subsequently, the membranes were blocked using 5% BSA (bovine serum albumin) for 1 h at room temperature and then hybridized sequentially using the primary antibodies and fluorescently labeled secondary antibodies. Signals were detected using the Odyssey infrared imaging system (Odyssey, LI-COR). The antibodies used are listed as follows: anti-BCL6 (1;1000, Cell Signaling Technology Cat# 14895, RRID:AB_2798638), anti-phospho-mTOR^S2448^ (1;1000, Cell Signaling Technology Cat# 2971, RRID:AB_330970), anti-mTOR (1;1000, Cell Signaling Technology Cat# 2972, RRID:AB_330978), anti-phospho-p70S6K^T389^ (1;1000, Cell Signaling Technology Cat# 9206, RRID:AB_2285392), anti-p70S6K (1;1000, Cell Signaling Technology Cat# 9202, RRID:AB_331676), anti-phospho-S6^S235/S236^ (1;1000, Cell Signaling Technology Cat# 2211, RRID:AB_331679), anti-S6 (1;1000, Cell Signaling Technology Cat# 2217, RRID:AB_331355), anti-phospho-γ-H2AX^S139^ (1;1000, Cell Signaling Technology Cat# 9718, RRID:AB_2118009), anti-PTEN (1;1000, Cell Signaling Technology Cat# 9559, RRID:AB_390810), anti-GAPDH (1;10000, Abcam Cat# ab181602, RRID:AB_2630358), anti-STAT1 (1;1000, Abclonal Cat# A19563, RRID:AB_2862669), and anti-IFNAR1 (1;1000, Proteintech Cat# 13083-1-AP, RRID:AB_2122626). The immunoblots are representative of three independent experiments.

### Enzyme-linked immunosorbent assay

To detect the cellular IFN-α and IFN-γ contents, cell lysates were extracted using RIPA buffer. The total protein concentrations were determined using BCA protein assay kit (Thermo Scientific), and IFN-α and IFN-γ protein concentrations were measured using a human IFN-α ELISA kit (1110012) and a human IFN-γ ELISA kit (1110002) from Dakewe Biotech, according to the manufacturer’s protocol.

### RNA sequencing

RNA-seq data were produced by Novogene (Beijing, China). Capan-2, H661, and PC9 cells were treated with dimethyl sulfoxide (DMSO) or etoposide at their respective IC_50_s for 24 h. Cells were harvested, and the total RNA was extracted using TRIzol reagent (Invitrogen) following the manufacturer’s protocol. A total of 1 μg RNA per sample was used as the input material for the RNA sample preparations. Libraries were prepared using the NEBNext UltraTM RNA Library Prep it for Illumina (NEB, USA) and library quality was assessed using the Agilent Bioanalyzer 2100 system. The clustering of the index-coded samples was performed using a cBot cluster generation system and a TruSeq PE cluster kit (Illumia) according to the manufacturer’s instructions. After cluster generation, the library preparations were sequenced on an Illumina platform and 150 bp paired-end reads were generated. Differential expression was analyzed using DESeq2 (RRID:SCR_000154). Pathway analysis was performed using gene set enrichment analysis (GSEA; http://software.broadinstitute.org/gsea/index.jsp).

### Single cell gel electrophoresis (comet) assay

Single cell electrophoresis (Neutral) was performed according to the manufacturer’s instructions (Trevigen). HCT116 and Capan-2 cells were treated with 10 μM etoposide, 10 μM BI3802, or both for 48 h. Afterward, cells were re-suspended in PBS at 2 × 10^5^ cells/mL and mixed with molten LMAgarose (at 37°C) at a ratio of 1:10. A 50 µL mixture was pipetted onto comet slides. The slides were solidified, and successively immersed in lysis solution and neutral electrophoresis buffer. The slides were then performed to electrophoresis, placed in a DNA precipitation solution, and stained using diluted SYBR® Gold. Signals were captured using a fluorescence microscope. DNA damage was quantified for 50 cells using the mean for each experimental condition, which was obtained by using Comet Score (TriTek) software.

### Animal experiments

For the human cancer cell xenograft mouse model, 6-week-old male BALB/cA nude mice were purchased from the National Rodent Laboratory Animal Resources (Shanghai, China). HCT116 cells (3 million per mouse) were injected subcutaneously into the flanks of nude mice. The primary *KRAS*-mutant lung cancer xenograft mouse model (LACPDX) was established as previously described (J. Wang et al., 2016). The patient-derived tumor tissues were cut into ∼15 mm^3^ fragments and implanted subcutaneously into BALB/cA nude mice using a trocar needle. For these two different xenograft mouse models, the tumors were measured using electronic calipers every other day, and the body was measured in parallel. When the tumor volume reached approximately 100 - 200 mm^3^, mice were randomized and treated with vehicle (dissolved in sterile water supplied with 0.5% CMC-Na), etoposide (10 mg/kg, orally, dissolved in corn oil), FX1 (5 mg/kg, intraperitoneally, dissolved in sterile water supplied with 0.5% CMC-Na) or etoposide + FX1. The tumor volumes were calculated using the formula, volume=length×width^2^×0.52. On day 16 or 24, the mice were sacrificed, and tumor tissues were excised, weighed and snap-frozen in liquid nitrogen for qPCR analysis, Western blotting analysis, and biochemistry testing. All animal experiments were conducted following a protocol approved by the East China Normal University Animal Care Committee.

### Statistical analysis

The data are presented as the mean ± S.E.M. unless otherwise stated. Statistical tests were performed using Microsoft Excel and GraphPad Prism Software version 8.0. For comparisons of two groups, a two-tailed unpaired *t*-test was used. For comparisons of multiple groups, one-way analysis of variance was used. Significance levels were set at * *P* < 0.05, ** *P* < 0.01, *** *P* < 0.001. Other specific tests applied are included in figure legends.

## Acknowledgements

We thank Dr. Yihua Chen (East China Normal University, Shanghai, China) for synthesizing and providing BCL6 inhibitors. We also thank Dr. Boyun Tang (Baygene Biotechnology, Shanghai, China) for RNA-seq data analysis.

## Availability of supporting data

RNA-seq data sets and the processed data that support the findings of this study have been deposited to the Gene Expression Omnibus (GEO) under accession ID: GSE161803.

## Authors’ Contributions

Y. L. designed and performed experiments, analyzed data and wrote the manuscript. J.F., performed experiments and revised the manuscript. K.Y., Y.L., K.L. performed and assisted with experiments. J.G., J.C. C.M. provided experimental supports and revised the manuscript. X.P. led the project, analyzed data and wrote the manuscript.

## Ethics Approval and Consent to participate

This study was approved by the Ethics Committee of the East China Normal University.

---

**Figure 1-figure supplement 1.**
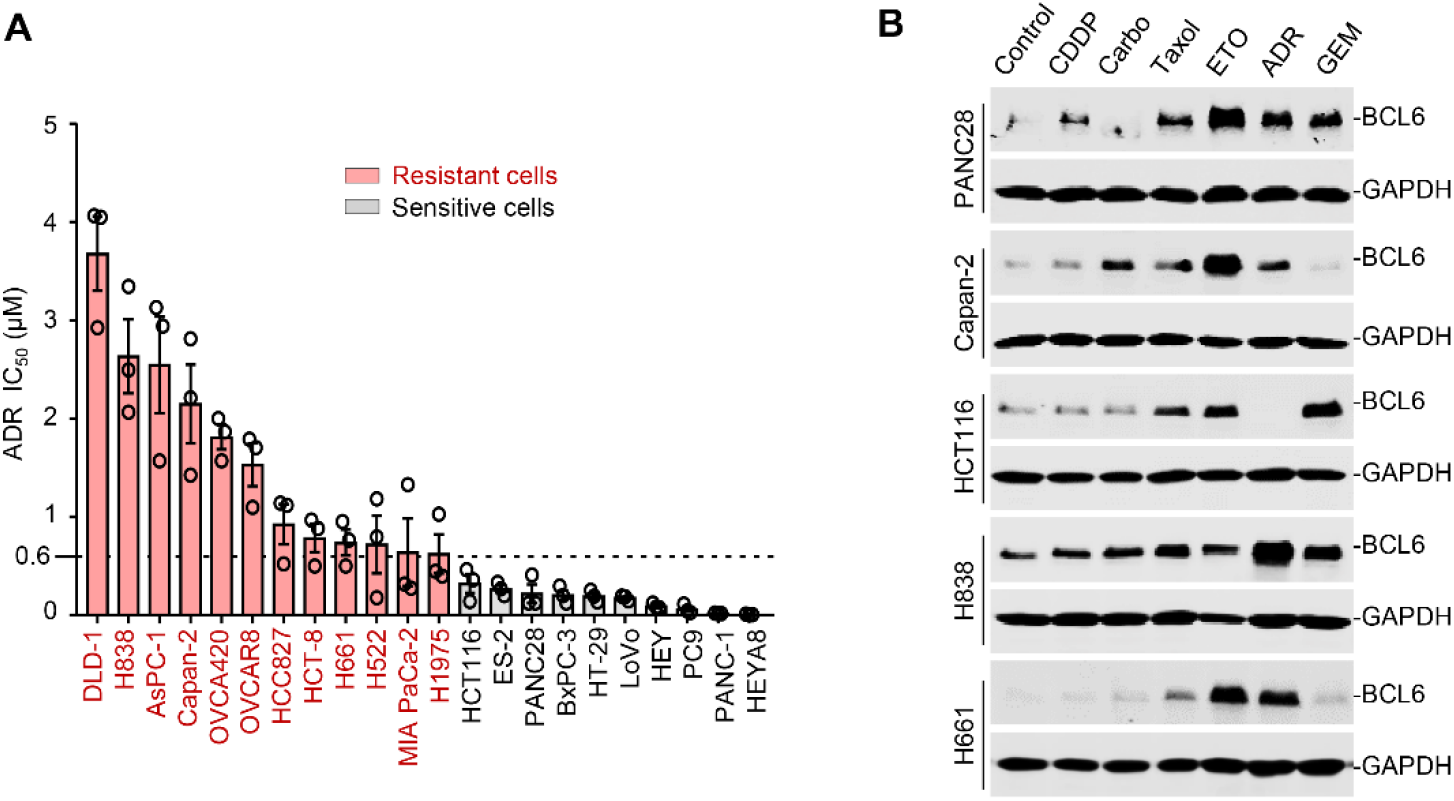
Genotoxic agents promote BCL6 expression. (**A**) Cell sensitivity to doxorubicin (ADR). Various cancer cell lines were treated with ADR at gradient concentrations for 48 h. IC_50_s were measured using SRB assays. Values are expressed as mean ± SEM of three independent experiments with triplates. ADR-resistant cell lines are marked in red. (**B**) BCL6 protein expression levels in different cancer cell lines in response to genotoxic agents. Cells were treated with indicated genotoxic agents at their respective IC_50_s for 24 h. Proteins lysates from each cell line were blotted individually. CDDP, cisplatin; Carbo, carboplatin; ETO, etoposide; GEM, gemcitabine.

**Figure 2-figure supplement 1.**
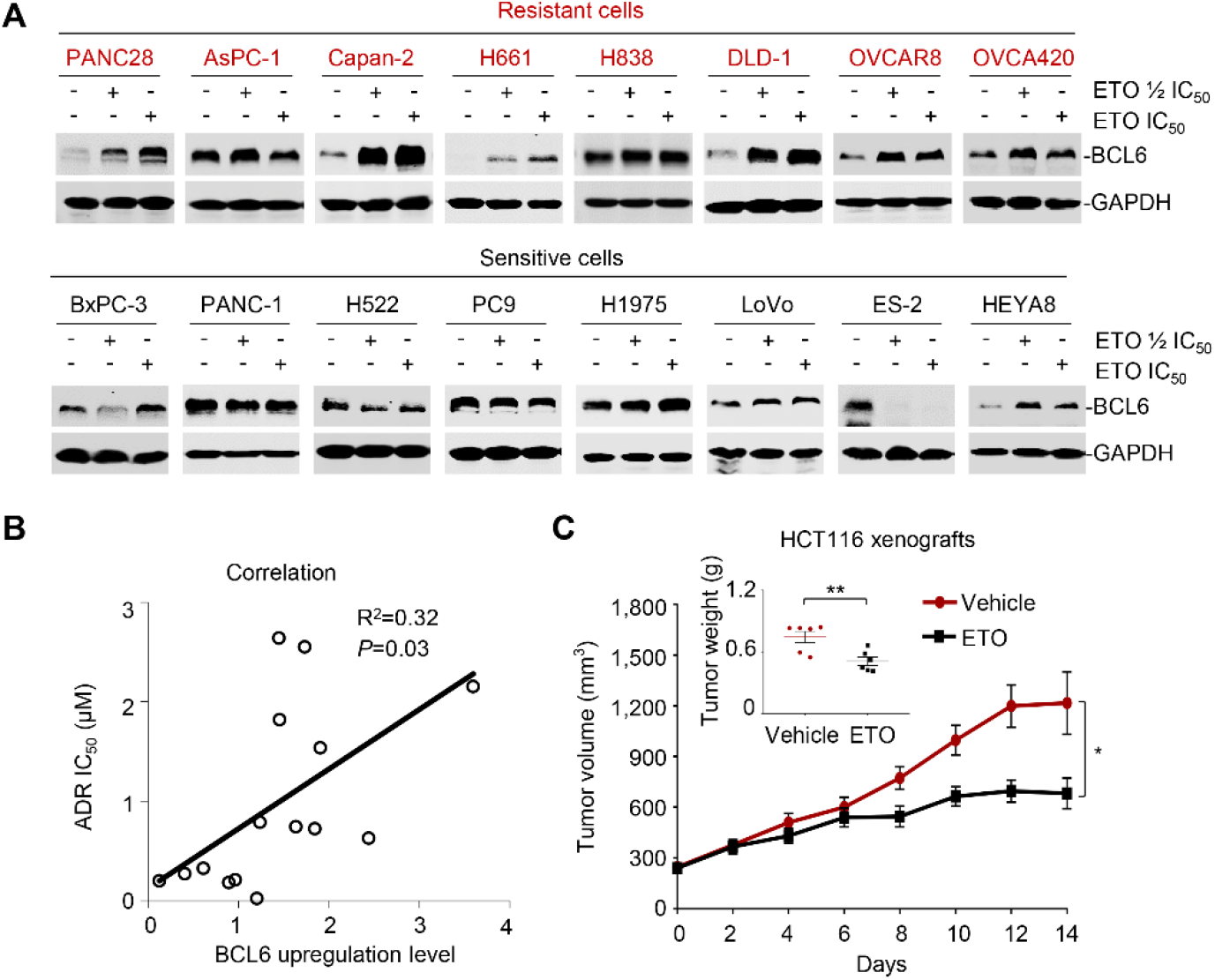
BCL6 upregulation is associated with therapy resistance. (**A**) ETO induced BCL6 protein expression. ETO-resistant or -sensitive cancer cells were treated with etoposide at their 1/2 IC_50_s or IC_50_s for 24 h, respectively. Proteins lysates from each cell line were blotted individually. (**B**) Correlation between BCL6 upregulation levels and ADR IC_50_s in various cancer cell lines. (**C**) Tumor volume curves and mean tumor weight on day 14. Data are expressed as mean ± SEM. \**P* < 0.05, \*\**P* < 0.01, unpaired, two tailed *t*-test, *n* = 6.

**Figure 3-figure supplement 1.**
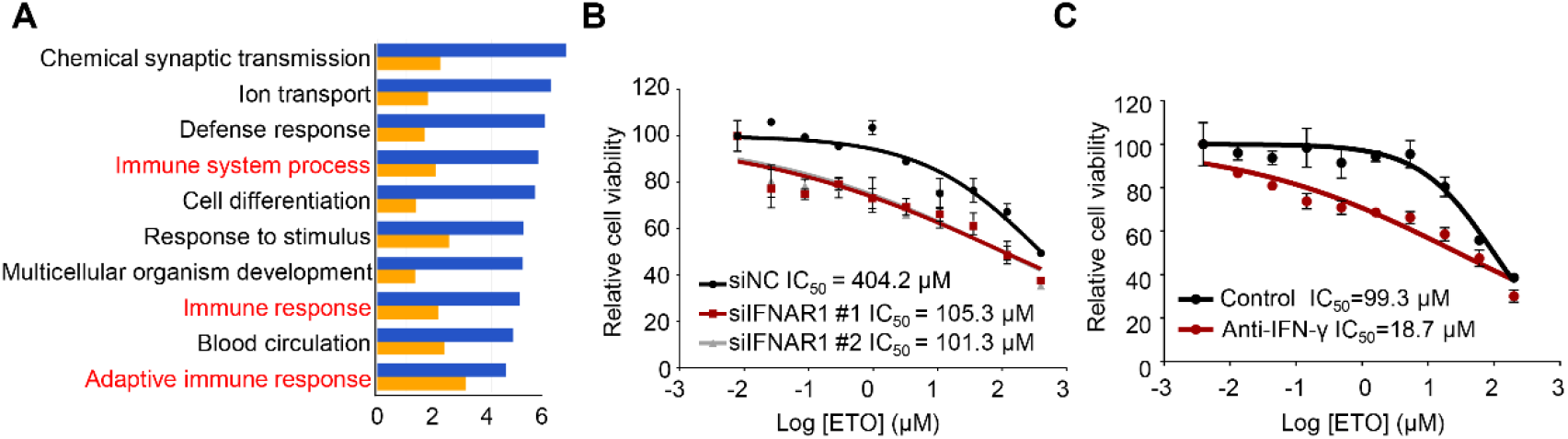
Genotoxic stress activates interferon responses. (**A**) GO analysis of RNA-seq data (ETO treatment group versus the control group). The top ten upregulated pathways in Capan-2, as indicated (*n* = 3). Graph displays category scores as log_10_ (P value) from Fisher’s exact test. (**B**) Silencing of IFNAR1 enhanced Capan-2 cells sensitivity to etoposide. Capan-2 cells were transfected with IFNAR1 siRNAs or the control siRNA for 48 h. Transfected cells were then exposed to etoposide at gradient concentrations for 48 h. Cell viability was detected using SRB assays. Values are expressed as mean ± SEM of two independent experiments. (**C**) Anti-IFN-ɣ antibody increased etoposide cytotoxicity. PANC28 cells were treated with etoposide at gradient concentrations for 48 h in the presence or absence of 10 μg/mL anti-IFN-ɣ antibody. Cell viability was detected using SRB assays. Values are expressed as mean ± SEM of three independent experiments.

**Figure 4-figure supplement 1.**
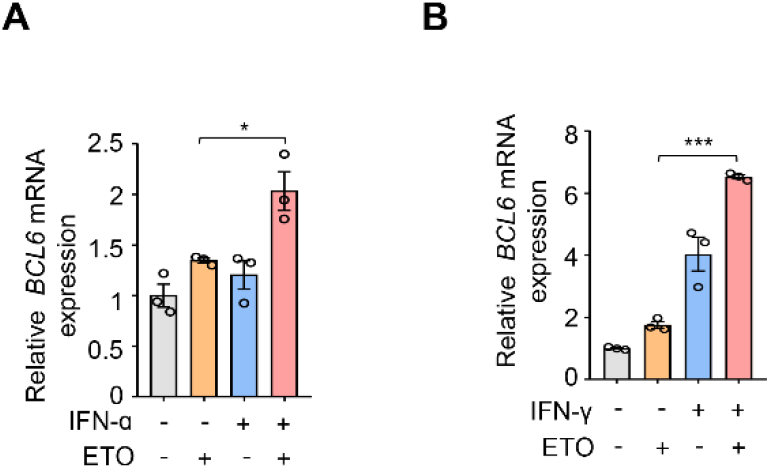
The interferon/STAT1 axis directly regulates BCL6 expression. **(A** and **B)** Relative BCL6 expression. Capan-2 cells were treated with 50 ng/mL IFN-ɑ (**A**) or 10 ng/mL IFN-γ (**B**) in the presence or absence of 50 μM etoposide. BCL6 mRNA levels were detected by qPCR assays. Values are expressed as mean ± SEM of three independent experiments. \**P* < 0.05, \*\*\**P* < 0.001, unpaired, two tailed *t*-test.

**Figure 6-figure supplement 1.**
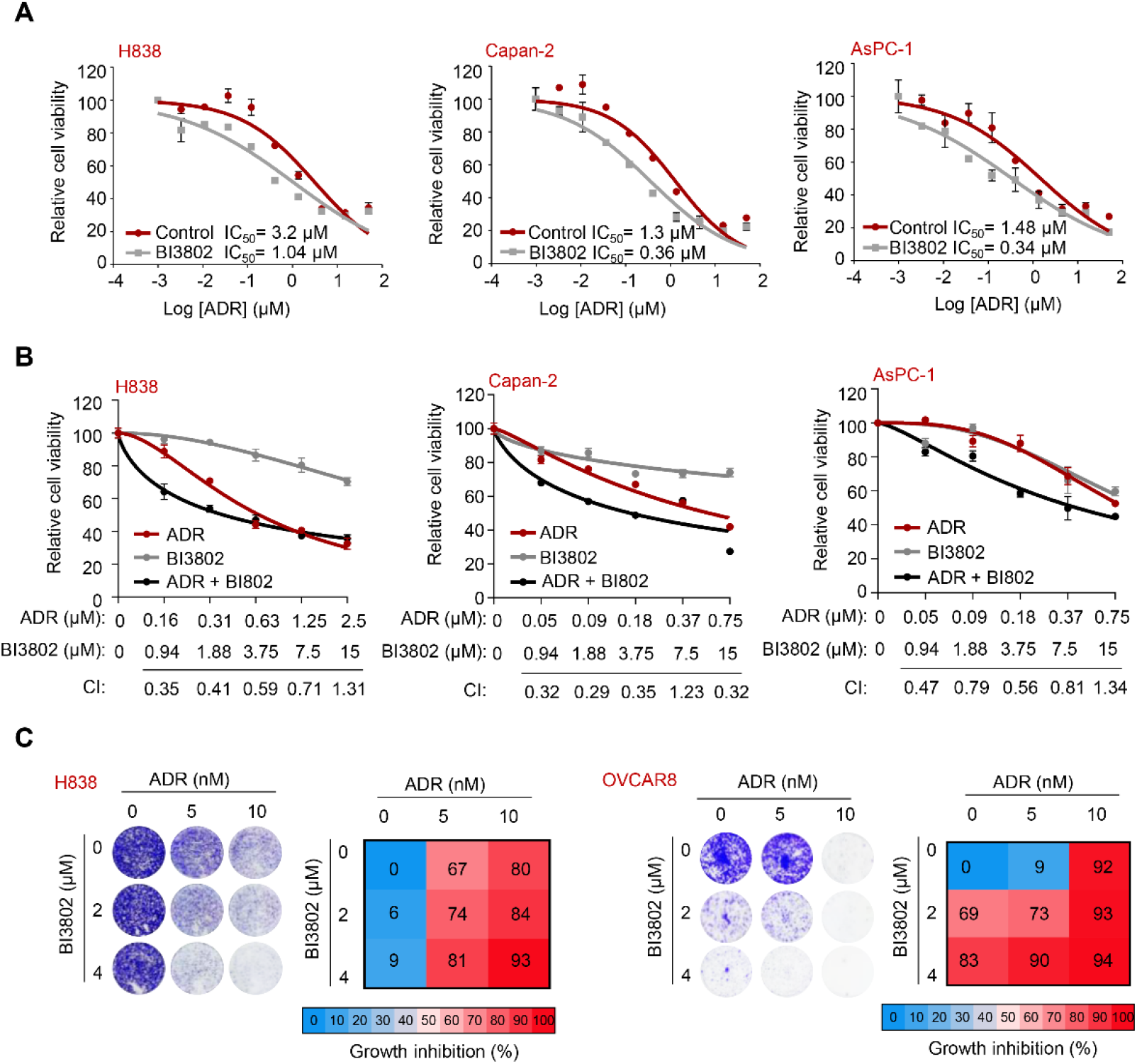
BCL6 inhibition sensitizes cancer cells to doxorubicin. (**A**) Increased sensitivity of cancer cells to doxorubicin. ADR-resistant cancer cells were treated with doxorubicin at gradient concentrations for 48 h in the presence of 10 μM BI3802. IC_50_s were measured using SRB assays. Values are expressed as mean ± SEM of two independent experiments. ADR-resistant cell lines are marked in red. (**B**) Cell viability of ADR-resistant cancer cells treated with different concentrations of doxorubicin in the combination with BI3802. Growth inhibition for three independent biological replicate experiments was averaged and input into CalcuSyn software to extrapolate CI values. CI values < 1 represent synergism. Values are expressed as mean ± SEM of three independent experiments. ADR-resistant cell lines are marked in red. (**C**) Representative long-term clonogenic assays (*left*) and quantified clonogenic growth inhibition data (*right*) for H838 and OVCAR8 cells treated with ADR, BI3802, or their combinations. Data are presented as mean of three independent experiments.

